# Tail Length and E525K Dilated Cardiomyopathy Mutant Alter Human β-Cardiac Myosin Super-Relaxed State

**DOI:** 10.1101/2023.12.07.570656

**Authors:** Sebastian Duno-Miranda, Shane R. Nelson, David V. Rasicci, Skylar L.M. Bodt, Joseph A. Cirilo, Duha Vang, Sivaraj Sivaramakrishnan, Christopher M. Yengo, David M. Warshaw

## Abstract

Dilated cardiomyopathy (DCM) is characterized by impaired cardiac function due to myocardial hypo-contractility and is associated with point mutations in β-cardiac myosin, the molecular motor that powers cardiac contraction. Myocardial function can be modulated through sequestration of myosin motors into an auto-inhibited "super relaxed" state (SRX), which is further stabilized by a structural state known as the "Interacting Heads Motif" (IHM). Therefore, hypo-contractility of DCM myocardium may result from: 1) reduced function of individual myosin, and/or; 2) decreased myosin availability due to increased IHM/SRX stabilization. To define the molecular impact of an established DCM myosin mutation, E525K, we characterized the biochemical and mechanical activity of wild-type (WT) and E525K human β-cardiac myosin constructs that differed in the length of their coiled-coil tail, which dictates their ability to form the IHM/SRX state. Single-headed (S1) and a short-tailed, double-headed (2HEP) myosin constructs exhibited low (∼10%) IHM/SRX content, actin-activated ATPase activity of ∼5s^-1^ and fast velocities in unloaded motility assays (∼2000nm/s). Double-headed, longer-tailed (15HEP, 25HEP) constructs exhibited higher IHM/SRX content (∼90%), and reduced actomyosin ATPase (<1s^-1^) and velocity (∼800nm/s). A simple analytical model suggests that reduced velocities may be attributed to IHM/SRX-dependent sequestration of myosin heads. Interestingly, the E525K 15HEP and 25HEP mutants showed no apparent impact on velocity or actomyosin ATPase at low ionic strength. However, at higher ionic strength, the E525K mutation stabilized the IHM/SRX state. Therefore, the E525K-associated DCM human cardiac hypo-contractility may be attributable to reduced myosin head availability caused by enhanced IHM/SRX stability.

**Summary:** This research investigates the E525K mutation in human β-cardiac myosin, crucial for heart contraction, and its role in causing Dilated Cardiomyopathy (DCM). It demonstrates that the length of the myosin tail influences its self-inhibition, and the E525K mutation strengthens this effect, potentially reducing heart contractility in DCM.

## Introduction

Dilated cardiomyopathy (DCM) is characterized by left ventricular dilation and systolic dysfunction, resulting in reduced cardiac output^1^ and eventually heart failure. Approximately 35% of DCM cases are monogenic, with mutations to genes encoding the heart muscle’s cytoskeletal and contractile proteins^2^. One etiologic hypothesis for DCM-associated mutations of β-cardiac myosin (M2β), the molecular motor that powers the heart^3^, is that these mutations negatively impact M2β function, which leads to myocardial hypo-contractility. Therefore, defining the impact of DCM-associated M2β mutations on molecular motor function is the first step in therapeutic design.

Upon cardiac muscle activation, ventricular force and motion generation originate within the sarcomere, the most basic contractile unit of cardiac muscle^4^. The sarcomere is composed of myosin thick filaments from which the myosin heads emanate (Fig. 1a), capturing the chemical energy of adenosine triphosphate (ATP) hydrolysis and converting it into mechanical energy as the motor interacts cyclically with the actin thin filament^4^. Thus, peak systolic force is determined by the number of attached myosin motors at any point in time, each of which generates an ∼1-2 pN intrinsic force^5,6^. Until recently, it was assumed that upon muscle activation, all myosin heads were available to interact with the thin filament. However, in relaxed striated muscles, including cardiac muscle, myosin motors adopt one of two distinct biochemical states, an auto-inhibited, super-relaxed (SRX) state, incapable of binding to the thin filament, and a disordered-relaxed (DRX) state that can be recruited upon activation to generate force^4,7^ (Fig. 1b,c). Therefore, the SRX/DRX ratio would determine the number of available motors that can attach to the thin filament and generate force and motion^8^. Although the SRX state is a biochemical state that is inherent to each of the myosin’s two heads^9^, the SRX state can be further stabilized by intramolecular electrostatic interactions, whereby the two heads fold back and interact with each other and the myosin coiled-coil tail^9,10^ (Fig. 1b, 2); an asymmetric structure called the "Interacting Heads Motif" (IHM)^11^. The IHM was first observed in electron micrographs of single myosin molecules^12^, with its presence now confirmed by cryo-electron microscopy of isolated native cardiac thick filaments^13^ and thick filaments within intact sarcomeres^14^. The IHM and its potential stabilization of the auto-inhibited SRX state offers a mechanism by which the number of available myosin heads competent to generate force can be regulated.

**Figure 1.**
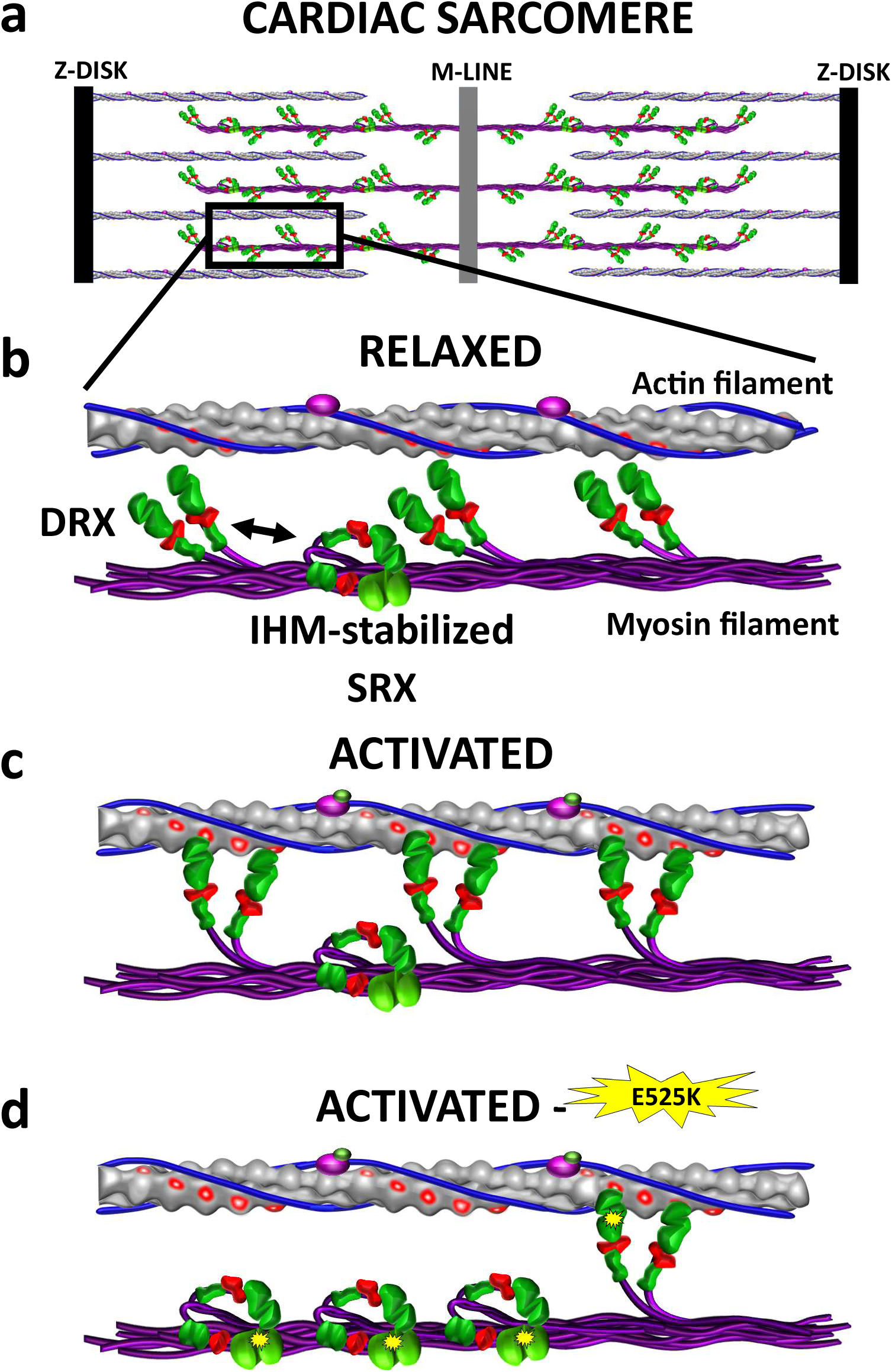
Cardiac myosin conformations and interactions with actin filaments in relaxed and activated Muscle. a) Representation of a cardiac sarcomere composed of interdigitated myosin filaments (purple) connected at the M-line and actin filaments anchored at the Z-disk. b) In relaxed muscle, myosin motors that emanate from the myosin filament backbone exist either in the enzymatically auto-inhibited, super-relaxed state (SRX), which can be stabilized by adopting the Interacting-Head-Motif (IHM) and precludes thin filament binding, or in the disordered-relaxed state (DRX) that is capable actin filament attachment and force generation upon cardiac activation. c) Once activated only DRX (i.e. active heads) myosin are recruited to generate force. d) Proposed mechanism by which the DCM-associated E525K M2β mutation (yellow starburst) stabilizes the IHM/SRX state, reducing the available pool of myosin motors for force generation, leading to cardiac hypo-contractility. Since most humans are heterozygote for cardiac myosin mutations, only one myosin head is depicted with the mutation.

**Figure 2.**
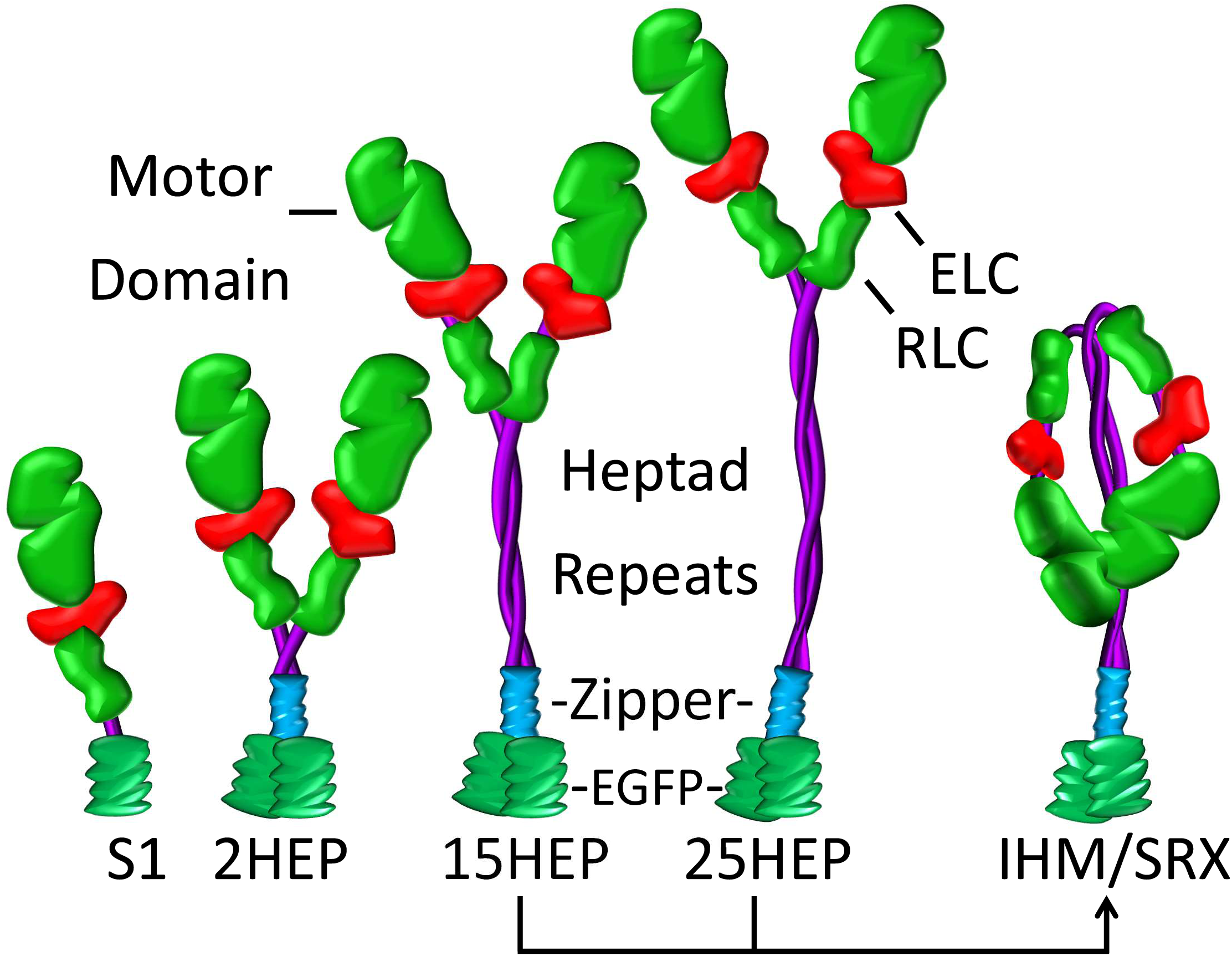
Various human single- and double-headed M2β constructs. Expressed human M2β constructs (single-headed: S1; double-headed with varying heptad (HEP) tail length: 2HEP, 15HEP, and 25HEP). All constructs have a regulatory light chain (RLC), essential light chain (ELC), and a C-terminal EGFP, with the double-headed constructs also containing a GCN4 leucine zipper to ensure dimerization. Only the 15HEP and 25HEP can adopt the IHM conformation, which presumably stabilizes the SRX state.

In contrast to DCM, hypertrophic cardiomyopathy (HCM) is also associated with M2β mutations, but is characterized by a hyper-contractile myocardium. HCM linked mutations have been shown to destabilize the IHM/SRX state, allowing more myosin heads to interact with the thin filament, offering a mechanistic explanation for the hyper-contractile HCM phenotype^15-17^. In fact, Mavacamten, an FDA approved, small-molecule therapeutic for HCM patients^18^, targets M2β and restores normal contractility by increasing the probability of the myosin IHM/SRX state. Accordingly, we hypothesize that the hypo-contractile phenotype observed with DCM-associated M2β mutations arises from enhanced stabilization of the IHM/SRX state, reducing the number of myosin motors available to generate the power required for normal cardiac output^8^ (Fig. 1d), although DCM-associated M2β mutations may also alter the motors’ intrinsic force and motion generation^19^. Therefore, in this study we chose to define the functional impact on the enzymatic and molecular mechanics of expressed human M2β harboring an established DCM-associated mutation (E525K), which localizes to an intramolecular contact region believed to stabilize the IHM^10,13^.

We build on our recently published biochemical and structural studies using single- and double-headed M2β constructs of both wildtype (WT) and the DCM-associated E525K mutant, which demonstrated that only the double-headed construct readily adopts the IHM/SRX state^10^. The present study provides biochemical and mechanical characterization of human M2β heavy meromyosin (HMM) constructs with varying lengths of coiled-coil domain (2, 15, and 25 heptad repeats, Fig. 2) to determine whether: 1) the presence of the IHM/SRX state in WT myosin affects *in vitro* actin filament motility, a model system for unloaded muscle shortening^20^, and; 2) the E525K DCM-associated mutation affects either the motor’s intrinsic enzymatic and mechanical performance and/or the capacity to adopt the IHM/SRX state. In fact, our new biochemical and *in vitro* motility data suggest that double-headed, longer-tailed (15 and 25 heptad) HMM constructs readily adopt the IHM/SRX state, leading to lower actin-activated ATPase activity and slower actin filament velocities compared to constructs incapable of adopting the IHM/SRX state. Surprisingly, in these longer-tailed HMM, the E525K DCM-associated mutation had little impact on the actin-activated ATPase activity and *in vitro* actin filament motility, under the low ionic strength conditions necessary to make these measurements. However, single ATP turnover studies that characterize the IHM/SRX state proportion over a range of ionic strengths, indicate that the E525K mutation does stabilize the IHM/SRX state and may be the dominant factor that reduces the number of available motors upon activation *in vivo*, potentially explaining the hypo-contractile DCM phenotype.

## Methods

### Protein expression and purification

The human M2β constructs were all derived from the *MYH7* gene (GenBank: AAA51837.1) (Fig. 2). As previously described^10^, the M2β HMM construct (amino acids 1-946), with 15 heptads of the coiled-coil (15HEP), is followed first by a GCN4 leucine zipper (sequence: MKQLEDKVEELLSKNYHLENEVARLKKLVGER), then a short linker (GSGKL), and a C-terminal series of EGFP, Avi-tag (GLNDIFEAQKIEWHE), and FLAG-tag (DYKDDDDK). HMM constructs with either a shorter 2 heptad coiled-coil (2HEP) (amino acids 1-855) or a longer 25 heptad coiled-coil (25HEP) (amino acids 1-1016) had the same C-terminal, leucine zipper, linker, EGFP, and Avi- and FLAG-tags. The M2β subfragment 1 construct (S1) (amino acids 1-842) had only the C-terminal EGFP, Avi- and FLAG-tags. The E525K mutation was introduced by Quikchange site-directed mutagenesis. The constructs were cloned into the pDual shuttle vector and the initial recombinant adenovirus stock was produced by Vector Biolabs (Malvern, PA) at a titer of 10^8^ plaque forming units per mL (pfu/mL). As previously described^21^, the virus was expanded by infection of Ad293 cells at a multiplicity of infection (MOI) of 3-5. The virus was harvested from the cells and purified by CsCl density sedimentation, giving a final virus titer of 10^10^-10^11^ pfu/mL.

The mouse skeletal muscle derived C2C12 cell line was used to express the various M2β constructs as described previously^22-24^. Briefly, C2C12 cells were grown to ∼90% confluency on tissue culture plates (145/20 mm) in growth media (DMEM with 9% FBS, 1% Penicillin-Streptomycin). On the day of infection, 20 plates were differentiated by changing the media to contain horse serum (DMEM with 9% horse serum, 1% FBS, 1% Penicillin-Streptomycin) and simultaneously infected with virus at 4 x 10^7^ pfu/ml. The cells were harvested for myosin purification 10-12 days after infection and M2β constructs were purified by FLAG affinity chromatography. At least 3 biological replicates of each construct were expressed and the same replicates were used for both biochemical and motility experiments described below. Actin was purified using acetone powder from rabbit skeletal muscle^25^.

### Steady–state Actin-activated ATPase assay

We utilized the NADH-coupled ATPase assay to examine the actin-activated ATPase activity at a range of actin concentrations (0 - 60 µM). Briefly, each construct (S1, 2HEP, 15HEP, and 25HEP) at 0.1 µM, was assayed in the presence of 1 mM ATP, and the NADH absorbance was monitored for a 200 second period in the stopped-flow apparatus (Applied Photophysics) at 25°C in MOPS 20 buffer (10 mM MOPS, pH 7.0, 20 mM KCl, 1 mM MgCl_2_, 1 mM EGTA, and 1 mM DTT). For the 0 µM actin condition, 1.0 µM of construct was used to improve signal to noise. The linear fit of the NADH absorbance as function of time was converted to the ADP produced per unit time using a standard curve with known ADP concentrations. The ATPase was expressed as the amount of product produced per second per molar concentration of myosin. The ATPase data plotted as a function of actin concentration was fit to the Michaelis-Menten equation to determine the maximal ATPase activity (k_cat_) and actin concentration at which ATPase is one-half maximal (K_ATPase_).

### Single turnover measurements

The fluorescence of 3’-O-(N-Methyl-anthraniloyl)-2’-deoxyadenosine-5’-triphosphate (mantATP) (Jena Biosciences) was measured with 290 nm excitation and a 395 nm long-pass emission filter in a stopped-flow apparatus (Applied Photophysics) at 25°C. Each myosin sample was incubated on ice for about 10 minutes in MOPS 20 buffer at varying KCl concentrations (20 - 150mM) prior to each experiment. Single mantATP turnover reactions were performed by incubating 0.25μM myosin constructs (S1, 2HEP, 15HEP, 25HEP) with 1μM mantATP for 30 seconds at room temperature. Subsequently, the complex was mixed with saturating ATP (2mM) in the stopped flow. Fluorescence decays were examined over a 1000 second period and were fitted to a two-exponential function to determine the SRX and DRX rate constants (Suppl. Fig. 1). Relative amplitudes of the fast and slow phase rate constants determined the fraction of myosin in the DRX and SRX states, respectively.

### *In vitro* motility assay

The movement of fluorescently-labeled actin filaments over an M2β-coated microscope slide surface was characterized using a modified version of previously established protocols^10,26,27^. In brief, the *in vitro* motility assay occurs within a sealed flow-through experimental chamber (i.e., flow-cell) constructed using a nitrocellulose-coated microscope coverslip. To minimize non-specific M2β surface interactions that may compromise the motor’s capacity to propel actin filaments, all M2β constructs were attached to the surface via an anti-GFP nanobody^28^. Preliminary *in vitro* motility data confirmed that the motion generating capacity of both the S1 and 2HEP constructs were compromised when the constructs were directly adhered non-specifically to the nitrocellulose surface (Suppl. Fig. 2). This was not the case for the longer-tailed 15HEP and 25HEP constructs. Regardless, all constructs were attached via the nanobody, which was diluted to 700nM in Actin Buffer (25mM KCl, 25mM Imidazole, 1mM EGTA, and 10mM DTT, pH 7.4) and infused into the flow-cell, incubated for 2 minutes, followed by a bovine serum albumin (BSA) wash (0.5 mg/mL BSA in Actin Buffer) to block any exposed nitrocellulose surface. Prior to M2β addition to the flow-cell, all constructs were subjected to a "dead-head" spindown to reduce the amount of non-functional M2β. Specifically, M2β was centrifuged in the presence of equimolar actin and 1 mM ATP in Myosin Buffer (300mM KCl, 25mM Imidazole, 1mM EGTA, 4mM MgCl_2_, 10mM DTT, pH 7.4) with the supernatant recovered for motility. M2β constructs were then infused into the flow-cell at concentrations ranging from 5-700nM in Myosin Buffer and incubated for 2 minutes. To eliminate any remaining non-functional myosin heads within the assay, unlabeled actin filaments (1μM in Actin Buffer) were infused into the flow-cell and incubated for 1 minute, after which an Actin Buffer wash with 1mM added MgATP released actin filaments from only the ATP-sensitive, functional myosin heads. This approach effectively removed damaged heads due to their ATP-insensitive and irreversible actin filament binding. Next, an Actin Buffer wash followed by infusion of 1-2.5nM vortexed (∼1s) Alexa-532-phalloidin-labeled actin filaments in Actin Buffer with 20% v/v O_2_ scavengers (0.1 μg/mL glucose oxidase, 0.018 μg/mL catalase, 2.3 μg/mL glucose) and then incubated for 1 minute. Finally, the flow-cell was washed extensively with Actin Buffer, with the final infusion of Motility Buffer (Actin Buffer with the addition of 1mM ATP, 0.5% methylcellulose, 2% v/v O_2_ scavengers, 1mM Creatine Phosphate, 1mg Creatine Phosphate Kinase), initiating actin filament motility. O_2_ scavengers were used to prevent oxidative damage, while methylcellulose was used to maintain actin filaments near the motility surface by reducing diffusional excursions.

For experiments where 2’-deoxyadenosine-5’-triphosphate (dATP) (GenScript, Piscataway, NJ) was used to limit the myosin SRX state^29^, dATP was substituted for ATP in all ATP-containing buffers. For M2β mixture experiments, the same protocol described above was used, with the exception that 25nM WT 2HEP was infused first, the flow-cell was then washed with Actin Buffer and after two minutes between 0nM and 125nM WT 15HEP was infused into the flow-cell and incubated for 2 minutes with the remaining steps the same. All motility experiments were imaged in Total Internal Reflection Fluorescence (TIRF) microscopy using a Nikon Eclipse TiU with Plan Apo objective (100x, 1.35 n.a.) and a Lumen 200W metal arc lamp (Prior Scientific) with actin filament images captured on a Turbo 620G camera (Stanford Photonics, Inc.) running Piper Controlled Imaging Desktop Software (Stanford Photonics, Inc). Individual motility movies of 30 seconds in length (10 FPS) were recorded at three different 45×60 micron regions of interest within a single flow-cell. All experiments were performed at 30°C.

### M2β motility surface occupancy

To assess the M2β motility surface occupancy, we took advantage of each myosin construct having an EGFP C-terminal tag (see above). M2β EGFP fluorescence in the flow-cell was imaged on the same TIRF microscope described above for the motility experiments. Therefore, each motility experiment had an accompanying estimate of the M2β surface occupancy. Three seconds long, M2β EGFP fluorescence movies (10 FPS) were taken at three different 45×60 micron regions of interest within the same flow-cell after recording motility. For each movie, we took the first 10 frames and created an average intensity z projection (ImageJ) and recorded its integrated total intensity. Then for the given flow-cell and myosin concentration, the total intensities for the three movies were averaged and used as a readout of EGFP fluorescence. The range of EGFP fluorescence intensity in arbitrary units associated with the different myosin concentrations was normalized to that obtained for the maximum 700nM myosin used. However, it was critical that the camera system gain be adjusted so that the EGFP fluorescence for the 700nM myosin concentration did not saturate the system’s fluorescence intensity readout. Once set, the same gain was used for flow-cells with lesser myosin concentrations.

### *In vitro* motility data analysis

For each motility movie that had 10-40 filaments per movie, depending on conditions, a mean actin filament velocity and fraction motile filaments were determined using a combination of the ImageJ plugin MTrack2 and Mean Squared Displacement (*MSD*) analysis. Briefly, *x,y* trajectories for individual actin filaments were obtained using MTrack2 and used to calculate *MSD* according to:

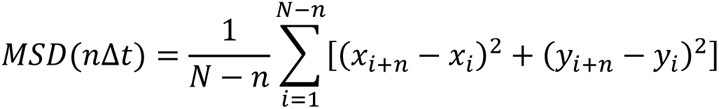

where *N* is the total number of frames in the trajectory, *n* is the number of frames for the time intervals over which the *MSD* is calculated (such that *n* ≤ *N*/4), *Δt* is the time between frames (100 ms), and *x_i_* and *y_i_* are the positions of the actin filament in frame *i*. As the relationship between MSD and time can be generalized as *MSD∼(nΔt)^α^*, we determined the diffusive exponent (*α*) as the slope of the relationship between *MSD* and the time interval (*nΔt*) on log-log axes. Filament trajectories of ≥ 20 data points that demonstrated directed motion (*α* > 1) were used to calculate a velocity from the slope of the relationship between √*MSD* and *nΔt*, while fraction of moving filaments was calculated as the fraction of all trajectories with *α*>1. For motility conditions where the myosin surface occupancy was varied, we then define a "Velocity Index" as the product of the mean velocity and the fraction of moving filaments at each surface occupancy. At least 3 biological replicates were characterized for all constructs and the replicate data for a given construct combined into mean velocity, mean fraction moving, and mean Velocity Index versus myosin surface occupancy (see Suppl. Fig. 3 for 2HEP WT example). Only the 2HEP WT Velocity Index versus myosin surface occupancy data were fit with a modified Hill dose-response equation using a Levenberg-Marquard algorithm (Origin(Pro) 2017, OriginLab Corporation, Northampton, MA, USA.), while all other construct relations were fit to predicted curves from an analytical model (see below).

### Statistics

Statistical comparisons between data sets were as follows. For both the motility surface EGFP fluorescence intensity versus myosin surface concentration and the Velocity Index versus myosin surface occupancy within WT groups or versus the E525K groups, data were compared using the non-parametric Kruskal-Wallis one-way analysis of variance. For experiments where the Velocity Index for constructs were compared at 100% myosin surface occupancy with and without the presence of dATP, the Student’s t-test was used for statistical comparisons. For both the Kruskal-Wallis and Student’s t-test a p value ≤0.05 was considered significantly different.

## Results

### WT M2β Steady-state ATPase activity

We previously studied the E525K DCM mutation and its impact on M2β structure and enzymatic activity^10^. However, we only compared the single-headed S1 and double-headed HMM 15HEP constructs. Here, we characterize these same constructs with both a shorter-tailed 2HEP and longer-tailed 25HEP as part of a more comprehensive study on how both the E525K mutation and M2β tail length impact biochemical and mechanical function.

The NADH-coupled ATPase assay was used to examine the steady-state actin-activated ATPase activity of each of our WT and E525K M2β constructs (S1, 2HEP, 15HEP, 25HEP; Fig. 2) in the absence and presence of actin (0, 5, 10, 20, 40, 60 µM) (Fig. 3). Since k_cat_ displayed considerable uncertainty in the long-tail constructs (15HEP and 25HEP), we compared the actin-activated ATPase at 60 µM actin to describe the relative differences between constructs and the impact of the mutation.

**Figure 3.**
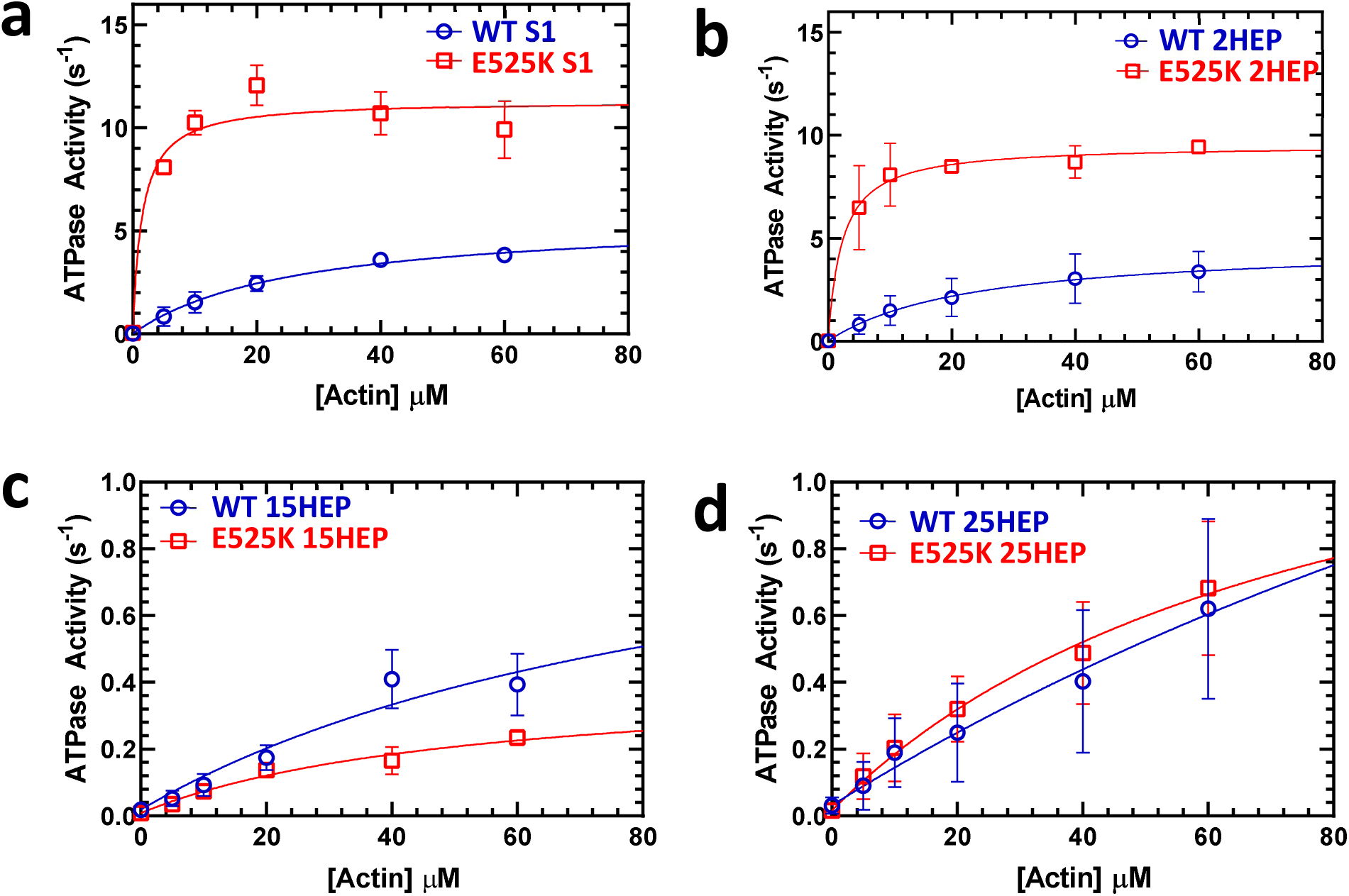
Steady-state ATPase activity of M2β constructs. a-d. Steady-state ATPase kinetics of WT and E525K M2β constructs were determined using an NADH-coupled assay across varying actin concentrations. E525K mutations in S1 and 2HEP constructs significantly increased actin-activated ATPase activity (∼3-fold) and decreased the actin concentration for half-maximal activity (K_ATPase_) by ∼18-fold. In contrast, 15HEP and 25HEP constructs showed reduced ATPase activities, with no significant difference between WT and E525K. Data are presented as mean ± SD of three experiments from separate protein preparations.

The actin-activated ATPase activity of the WT S1 and 15HEP HMM constructs containing an EGFP tag were similar to our previously published results without the EGFP tag^30^, suggesting that the C-terminal EGFP did not alter the ATPase activity (Table 1). As before, the WT 15HEP construct showed a ∼10-fold reduced actin-activated ATPase activity compared to S1 (Fig. 3a,c; Table 1), which we previously interpreted as evidence for the 15HEP construct adopting the IHM/SRX state^10,30^. Interestingly, the short-tailed, double-headed WT 2HEP construct displayed similar actin-activated ATPase activity to WT S1 (Fig. 3a,b), with similar *K*_ATPase_ and *k*_cat_ (Table 1). Therefore, on a per head basis the S1 and 2HEP were enzymatically indistinguishable (p>0.05). As with the 15HEP construct, the longer-tailed 25HEP construct displayed a reduced actin-activated ATPase activity compared to either the S1 or short-tailed 2HEP (Fig. 3d; Table 1), consistent with its ability to form the auto-inhibited, IHM/SRX state.

**Table 1.** Steady-state actin-activated ATPase measurements (± SE).

| Construct | $v_0$<br>($s^{-1}$) | ATPase @ 60<br>$\mu M$ actin ($s^{-1}$) | $K_{ATPase}$<br>($\mu M$) | $k_{cat}$<br>($s^{-1}$) |
| --- | --- | --- | --- | --- |
| WT S1, N=3 | 0.02±0.01 | 3.8±0.2 | 26.5±5.6 | 5.7±0.5 |
| WT 2HEP, N=4 | 0.03±0.03 | 3.4±1.0 | 23.4±11.5 | 4.7±1.0 |
| WT 15HEP, N=3 | 0.02±0.02 | 0.4±0.1 | 99.9±64.5 | 1.1±0.5 |
| WT 25HEP, N=4 | 0.03±0.02 | 0.6±0.3 | ND | ND |
| E525K S1, N=3 | 0.04±0.01 | 9.9±1.4** | 1.5±0.6* | 11.3±0.5** |
| E525K 2HEP, N=3 | 0.03±0.04 | 9.4±0.3** | 2.2±0.8* | 9.5±0.5* |
| E525K 15HEP, N=3 | 0.01±0.02 | 0.2±0.1 | 53.3±20.8 | 0.41±0.1 |
| E525K 25HEP, N=4 | 0.02±0.02 | 0.7±0.2 | ND | ND |

\*P<0.05 or \*\*P<0.005 for comparison of E525K and WT

### WT Single ATP turnover assays to measure the M2β SRX state population

The single ATP turnover assay is a powerful technique to understand the kinetics of a single round of ATP hydrolysis by a given enzyme. In the case of M2β in the absence of actin, the single ATP turnover assay has been particularly useful in examining the fraction of myosin molecules that populate the SRX state, as it hydrolyzes ATP 5-10 fold slower than the uninhibited myosin in the disordered-relaxed state (DRX). Therefore, we performed single ATP turnover assays to determine the SRX fraction in each of our WT and E525K M2β constructs (S1, 2HEP, 15HEP, 25HEP; Fig. 4, Suppl. Fig. 1).

**Figure 4.**
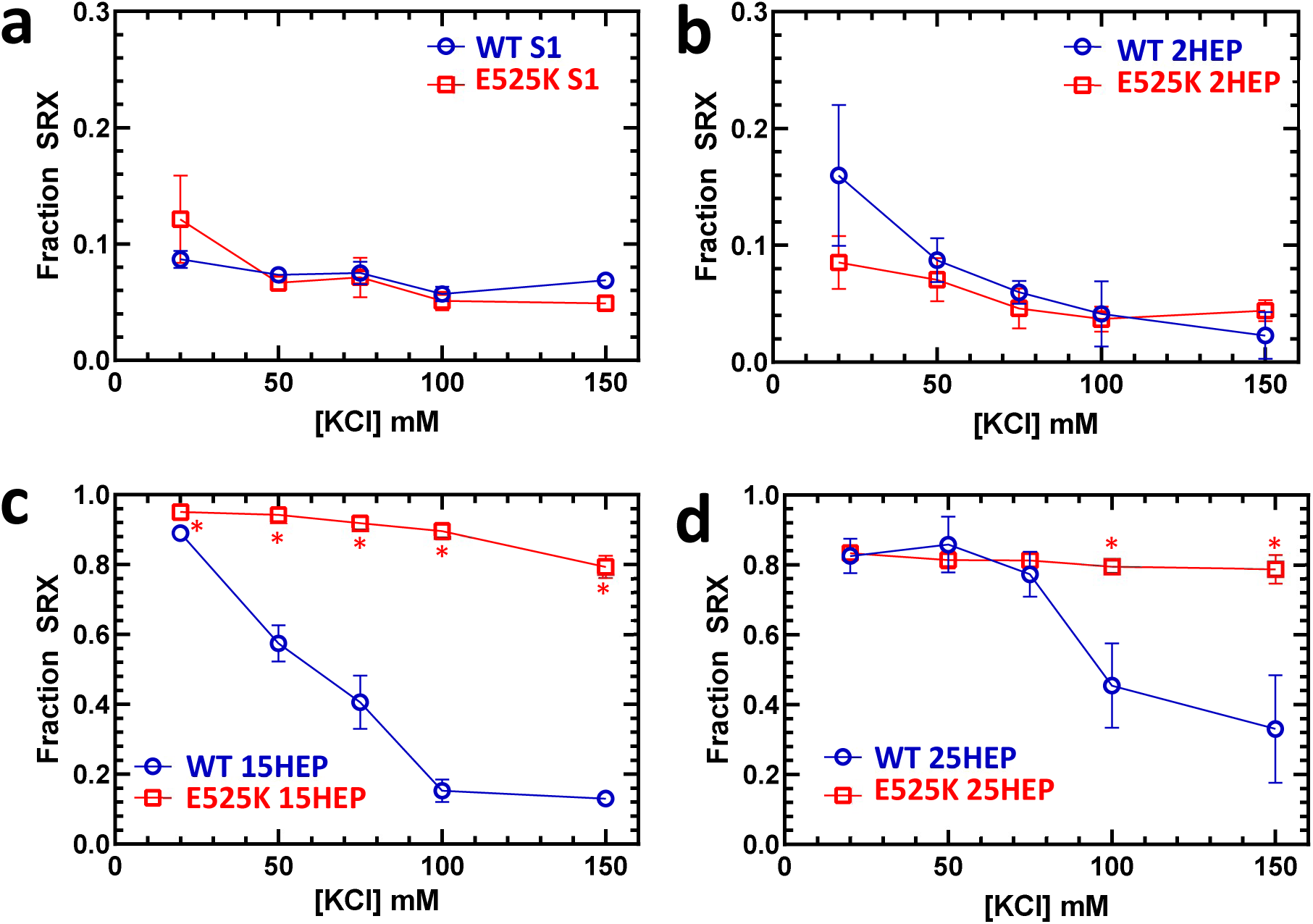
Assessment of IHM/SRX state fraction via single ATP turnover. a-d. Single turnover of mantATP by M2β constructs was analyzed at different KCl concentrations to deduce the fraction of myosin heads in the IHM/SRX state. WT and E525K constructs at 0.25 µM were pre-incubated with mantATP (1 µM) for approximately 30 seconds before introducing saturating ATP (2 mM). The fluorescence transients were fitted to a two-exponential decay function to infer the SRX state’s fraction and kinetics. Notably, long-tailed (15 & 25HEP; c & d) constructs displayed a greater SRX fraction than S1 or 2HEP (a & b), which was sensitive to salt concentration only in the WT constructs with significant differences identified by asterisks (*) (15HEP, p<0.005; 25HEP, p<0.05). The mean ± SD of three experiments from separate protein preparations is reported.

As in our previous study, single-headed WT S1 constructs exhibit an inherently low percentage of SRX heads regardless of the ionic strength (Fig. 4a; Tables 2,3), while the double-headed, long-tailed WT 15HEP construct populated the SRX state in an ionic strength-dependent manner, with nearly 90% SRX myosin heads at low salt (20 mM KCl) and 10% SRX at high salt (150 mM KCl) (Fig. 4c; Tables 2,3). These data provide evidence that the reduced steady state actin-activated ATPase for the 15HEP described above at low salt (Fig. 3c; Table 1) reflect electrostatic intramolecular interactions between the heads and long coiled-coil tail stabilizing the auto-inhibited IHM/SRX state. The tail length is critical to this inhibition, as the WT 2HEP construct showed a small SRX fraction (2-16%) at all KCl concentrations similar to the WT S1 (Fig. 4b; Tables 2,3). Interestingly, the longer-tailed WT 25 HEP construct demonstrated a high SRX content, but with a less pronounced ionic strength-dependence compared to the WT 15HEP, suggesting the additional 10 heptads of coiled-coil further stabilize the auto-inhibited state (Fig. 4d; Tables 2,3).

**Table 2.** Single mantATP turnover measurements for WT and E525K constructs (N=3, ± SD).

| KCl (mM) | DRX Rate<br>(s<sup>-1</sup>) | SRX Rate<br>(s<sup>-1</sup>) | SRX Fraction<br>(%) |
| --- | --- | --- | --- |
| WT S1 |  |  |  |
| 20 | 0.034±0.002 | 0.0020±0.0002 | 8.7±0.7 |
| 50 | 0.034±0.001 | 0.0029±0.0007 | 7.4±0.3 |
| 75 | 0.030±0.001 | 0.0016±0.0001 | 7.5±1.0 |
| 100 | 0.028±0.002 | 0.0029±0.0012 | 5.7±0.6 |
| 150 | 0.025±0.001 | 0.0028±0.0008 | 6.8±0.3 |
| E525K S1 |  |  |  |
| 20 | 0.072±0.005 | 0.0019±0.0005 | 12.1±3.8 |
| 50 | 0.064±0.003 | 0.0023±0.0003 | 6.6±0.3 |
| 75 | 0.063±0.005 | 0.0017±0.0004 | 7.1±1.7 |
| 100 | 0.064±0.006 | 0.0023±0.0008 | 5.1±0.8 |
| 150 | 0.061±0.001 | 0.0024±0.0005 | 4.9±0.2** |

P<0.05 for comparison of E525K and WT
| KCl (mM) | DRX Rate<br>(s<sup>-1</sup>) | SRX Rate<br>(s<sup>-1</sup>) | SRX Fraction<br>(%) |
| --- | --- | --- | --- |
| WT 2HEP |  |  |  |
| 20 | 0.022±0.001 | 0.0066±0.0017 | 16.0±6.0 |
| 50 | 0.025±0.001 | 0.0053±0.0006 | 8.7±1.9 |
| 75 | 0.028±0.001 | 0.0047±0.0031 | 6.0±1.0 |
| 100 | 0.030±0.001 | 0.0047±0.0027 | 4.1±2.8 |
| 150 | 0.031±0.002 | 0.0038±0.0036 | 2.3±2.0 |
| E525K 2HEP |  |  |  |
| 20 | 0.045±0.002 | 0.0033±0.0005 | 8.5±2.3 |
| 50 | 0.059±0.002 | 0.0073±0.0030 | 7.1±1.9 |
| 75 | 0.063±0.003 | 0.0052±0.0013 | 4.6±1.7 |
| 100 | 0.062±0.001 | 0.0040±0.0018 | 3.7±1.1 |
| 150 | 0.068±0.001 | 0.0042±0.0046 | 4.4±0.9 |
| WT 15HEP |  |  |  |
| 20 | 0.014±0.002 | 0.0033±0.0001 | 89.1±0.8 |
| 50 | 0.013±0.001 | 0.0054±0.0005 | 57.4±5.2 |
| 75 | 0.015±0.002 | 0.0069±0.0009 | 40.6±7.6 |
| 100 | 0.016±0.001 | 0.0058±0.0004 | 15.3±3.3 |
| 150 | 0.018±0.001 | 0.0071±0.0008 | 13.0±1.7 |
| E525K 15HEP |  |  |  |
| 20 | 0.027±0.008 | 0.0022±0.0002 | 95.0±0.7** |
| 50 | 0.019±0.002 | 0.0026±0.0001 | 94.2±0.7** |
| 75 | 0.016±0.006 | 0.0030±0.0001 | 91.8±1.6** |
| 100 | 0.014±0.003 | 0.0033±0.0001 | 89.6±2.3** |
| 150 | 0.013±0.002 | 0.0042±0.0001 | 79.4±3.2** |

\*\*P<0.005 for comparison of E525K and WT

| KCl (mM) | DRX Rate<br>(s<sup>-1</sup>) | SRX Rate<br>(s<sup>-1</sup>) | SRX Fraction<br>(%) |
| --- | --- | --- | --- |
| WT 25HEP |  |  |  |
| 20 | 0.036±0.009 | 0.0030±0.0001 | 82.5±5.0 |
| 50 | 0.033±0.016 | 0.0039±0.0004 | 85.8±8.0 |
| 75 | 0.025±0.012 | 0.0039±0.0004 | 77.3±6.4 |
| 100 | 0.011±0.003 | 0.0029±0.0005 | 45.4±12.1 |
| 150 | 0.010±0.001 | 0.0032±0.0008 | 33.1±15.4 |
| E525K 25HEP |  |  |  |
| 20 | 0.029±0.003 | 0.0030±0.0001 | 83.3±0.2 |
| 50 | 0.027±0.006 | 0.0034±0.0002 | 81.4±0.2 |
| 75 | 0.027±0.004 | 0.0041±0.0006 | 81.3±1.4 |
| 100 | 0.034±0.003 | 0.0048±0.0002 | 79.5±1.4* |
| 150 | 0.039±0.011 | 0.0061±0.0002 | 78.8±4.0* |

\*P<0.05 for comparison of E525K and WT

**Table 3.**
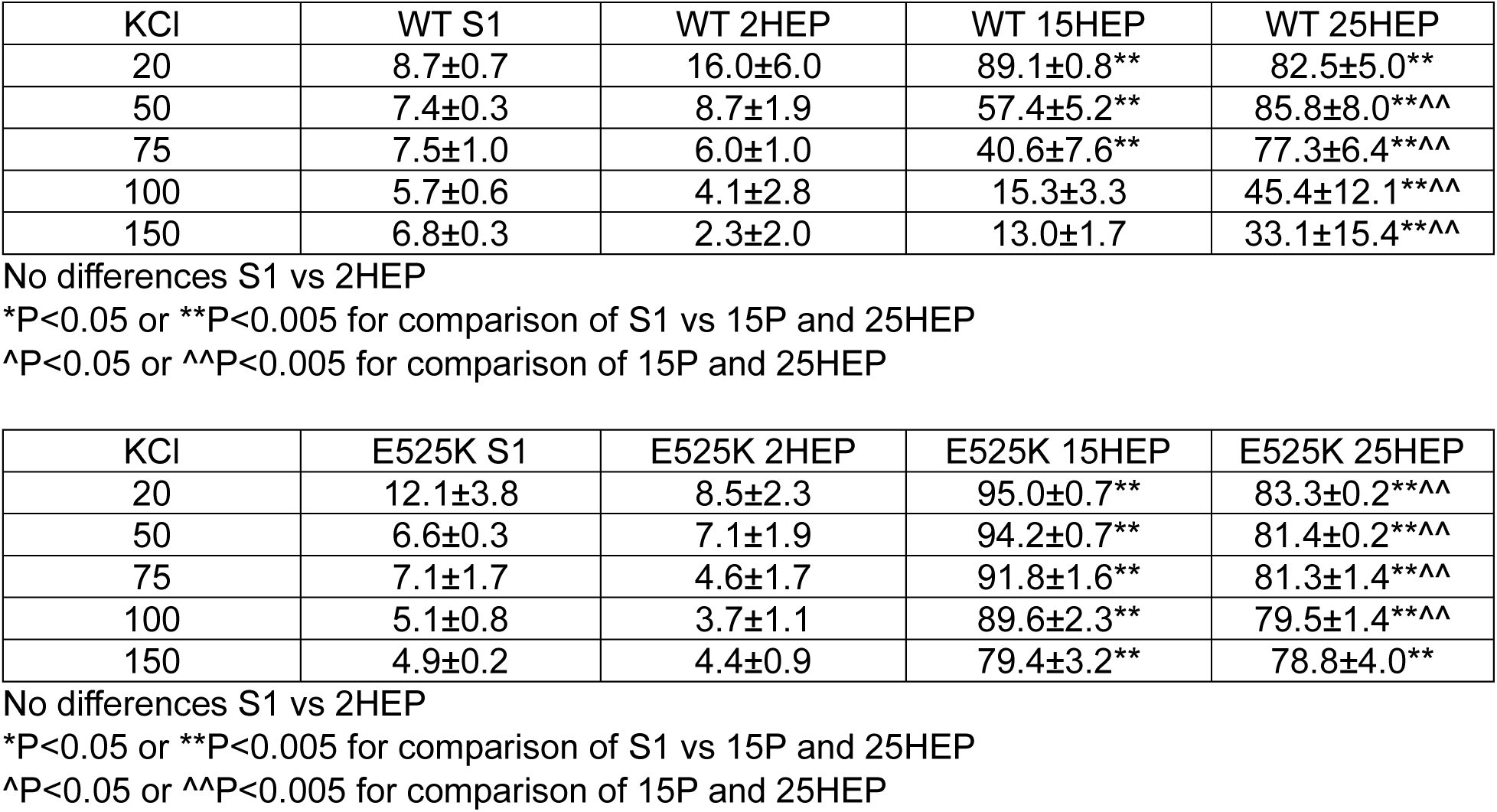
Comparison of SRX percentages for all M2β constructs at varying salt concentration.

### Defining M2β surface occupancy in the motility assay

Knowing that the various M2β constructs can adopt an IHM/SRX state, do these auto-inhibited motors impact *in vitro* motility? If so, then it is critical that the total motor number on the motility surface be estimated, which would include both the active and auto-inhibited (IHM/SRX) motors. Therefore, the C-terminal EGFP tag on each construct served both as a fluorescence readout for motor counting, as well as an attachment strategy to the motility surface via an anti-GFP nanobody (see Methods). All WT and E525K mutant constructs were introduced into the flow-cell at various concentrations (0-700nM). With increasing M2β concentration, the overall fluorescence intensity increased (Fig. 5a) and was well fit with a hyperbolic saturation binding model (Fig. 5b). The apparent surface affinity as characterized by an averaged K_app_ of 147 ± 33 nM for the WT double-headed constructs (2HEP, 15HEP, 25HEP) was no different than the corresponding E525K mutants with an averaged K_app_ of 165 ± 22 nM, suggesting that neither the tail length, nor E525K mutation impact the binding to the nanobody (p = 0.99). As expected, given that S1 has a single EGFP tag, the absolute fluorescence was lower at all WT S1 concentrations when compared to the double-headed WT 2HEP (Fig. 5b). Accordingly, WT S1 surface binding showed a lower K_app_ of 93 ± 27 nM.

**Figure 5.**
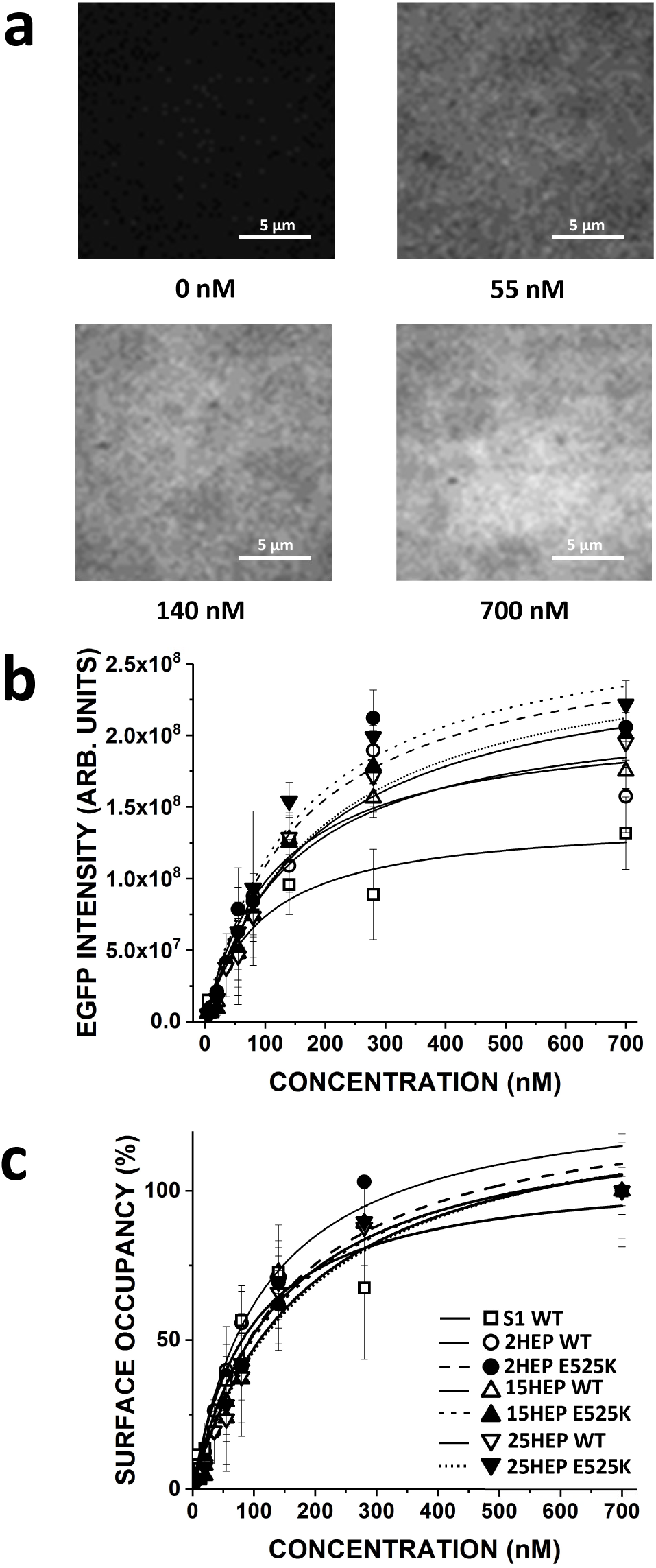
M2β construct motility assay surface occupancy estimation. a) M2β constructs were attached to the motility assay glass surface via their C-terminal EGFP to preadhered anti-GFP nanobodies. Increasing M2β binding showed increasing surface EGFP fluorescence intensity. b) Surface EGFP fluorescence intensity increased as a function of M2β construct concentration infused into motility assay flowcell. Fluorescence intensity followed saturable binding behavior with an apparent dissociation constant, *K_app_* (nM): *K*_app_S1 WT__ = 93 ± 27, *K*_app_2HEP WT__ = 116 ± 42, *K*_app_15HEP WT__ = 143 ± 32, *K*_app_25HEP WT__ = 181 ± 36, *K*_app_2HEP E525K__ = 152 ± 38, *K*_app_15HEP E525K__ = 191 ± 45, *K*_app_25HEP E525K__ = 153 ± 34. No difference in the surface EGPF intensity across WT and E525K mutants (p = 0.98). c) Data in “b” normalized to fluorescence at 700 nM M2β concentration and converted to Surface Occupancy. Data are mean ± SD of at least three experiments from separate protein preparations.

Once defined, each surface fluorescence curve was normalized to the fluorescence intensity at the 700 nM construct loading concentration and converted to percent surface occupancy (Fig. 5c). This allowed *in vitro* motility data across all constructs to be compared as a function of myosin surface occupancy regardless of the functional capacity of the construct (i.e., active versus auto-inhibited IHM/SRX).

### WT M2β *in vitro* motility

The *in vitro* motility for the various WT constructs was characterized by a Velocity Index (see Methods, Fig. 6), which is the product of the motile fraction of filaments and their velocity, providing a better metric of activity for very low densities of myosin. For limiting densities of low duty ratio (∼5%) motors, such as muscle myosin^31^, actin velocity is limited by the rate of cross-bridge attachment. Thus, actin velocity increases with the number of motors interacting with the actin filament (Suppl. Fig. 3a), saturating when at least one motor contributes to actin filament movement at any point in time. However, if additional motors simultaneously interact with the actin filament, no further increase in velocity is observed^32^. Therefore, if changes in the IHM/SRX state occur due either to the construct tail length or E525K mutation, which could alter the effective number of available motors, then comparing maximum velocities alone would be an insensitive measure. Therefore, changes in Velocity Index across a range of myosin surface occupancies will allow effective alterations in motor number due to construct differences or mutation to be identified.

**Figure 6.**
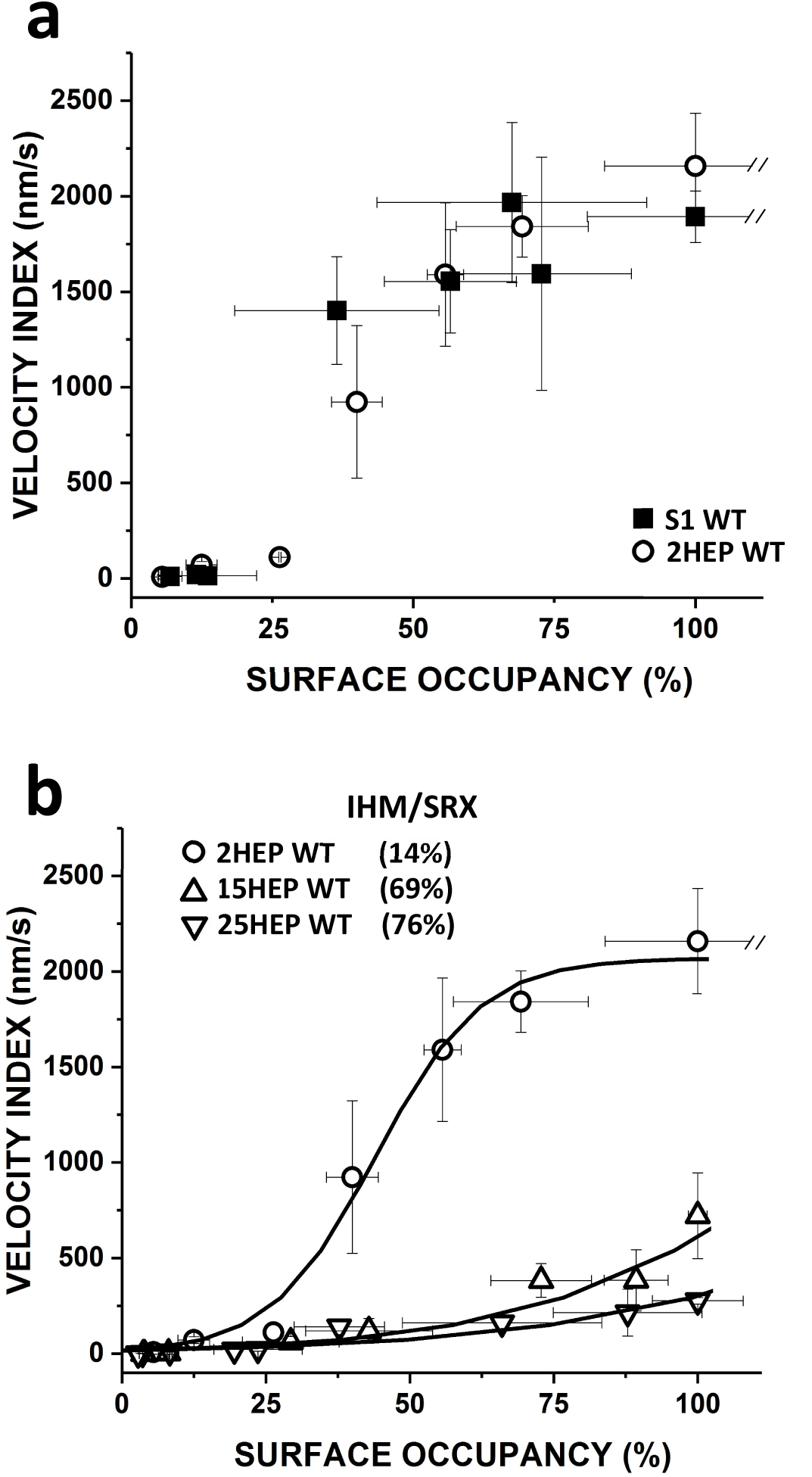
Motility analysis of WT human M2β constructs. a) The Velocity Index versus Surface Occupancy for WT S1 and 2HEP constructs showed a dose-response-like characteristic that saturated at ∼2,000nm/s and were indistinguishable (p = 0.87). b) The Velocity Index versus Surface Occupancy for WT 2HEP, 15HEP, and 25HEP constructs. The longer-tailed WT 15HEP and 25HEP constructs had indistinguishable Velocity Indices (p = 0.87), but slower at each myosin surface occupancy compared to WT 2HEP. The WT 2HEP Velocity Index versus Surface Occupancy data were fitted to a Hill dose-response relationship (solid curve) and then used as the basis for an analytical model (see Results) to predict the IHM/SRX content for each construct. Predicted IHM/SRX percentages (legend in figure) and associated fitted curve (solid curve) are shown. All *in vitro* motility experiments were performed in low salt (25 mM KCl). Data points are presented as mean ± SD of at least three experiments from separate protein preparations.

The Velocity Index relationships for the WT S1 and 2HEP constructs showed a dose-response-like characteristic that saturated at ∼2,000nm/s (Fig. 6a) and were indistinguishable (p = 0.87). The increasing Velocity Index with myosin surface occupancy suggests that motor number was rate limiting, which is supported by the fraction moving filaments being low at the low myosin surface occupancies (see Suppl. Fig. 3b). The similarity in the Velocity Indices for the WT S1 and 2HEP constructs may reflect the similarity in their steady state actin-activated and single turnover ATPase measurements (Figs. 3,4). Although the longer-tailed WT 15HEP and 25HEP constructs had Velocity Indices which were not different from each other (p = 0.87) (Fig. 6b), their Velocity Indices were slower at each myosin surface occupancy compared to the WT S1 and 2HEP constructs, being >60% slower at 100% myosin surface occupancy (Fig. 6a,b).

### Modeling the effect of the IHM/SRX on *in vitro* motility

With reductions in the actin-activated ATPase activities of the long-tailed WT 15HEP and 25HEP (Fig. 3c,d) compared to the WT S1 and 2HEP (Fig. 3a,b) constructs being attributed to the presence of IHM/SRX heads (Fig. 4), we developed a simple analytical model whereby the presence of IHM/SRX myosin for the long-tailed constructs accounts for a reduction in the Velocity Index as a function of myosin surface occupancy (Fig. 6). The model (see Supplemental Methods for model fitting details) was based on the following assumptions:

1. The myosin surface occupancy reflects the total number of myosin motors on the surface (Fig. 5);
2. An IHM/SRX motor does not generate or impede actin movement, i.e., it is mechanically silent, thus effectively reducing the number of active heads on the motility surface;
3. Each construct’s proportion of active and IHM/SRX is maintained at all myosin surface occupancies;
4. the Velocity Index versus myosin surface occupancy of the WT 2HEP construct (Fig. 6a) reflects a population of motors having 84% active and 16% IHM/SRX heads (Fig. 4b; Table 2);
5. As they share the same motor domain, the maximum theoretical Velocity Index for all constructs is that experimentally observed for saturating WT 2HEP (2,300nm/s) (Fig. 6).

Based on these assumptions, the WT 2HEP Velocity Index data were fitted to a modified Hill dose-response relationship and then used as a baseline curve for a construct that has 16% IHM/SRX content (Figs. 4b, 6b; Table 2). Using this same curve, a family of curves was generated with the only free parameter being the percent IHM/SRX, which was varied between 0% and 100% IHM/SRX (Suppl. Methods and Suppl. Fig. 4b). Using this family of curves, the best least squared error fits to the WT 2HEP, 15HEP, and 25HEP Velocity Index data predict IHM/SRX percentages of 14±13%, 69±6%, and 76±4%, respectively (Fig. 6b, solid curves; Suppl. Methods and Suppl. Fig. 4 for fitting and estimate error determination), in close agreement with results from single turnover experiments (Fig. 4).

To test the predictive nature of our model and its underlying assumptions, most importantly that each construct has a consistent proportion of IHM/SRX content, we performed a mixtures motility assay by adhering a constant 25nM WT 2HEP (14% IHM/SRX) with an additional 0nM to 125nM WT 15HEP (69% IHM/SRX). The resultant Velocity Index versus myosin surface occupancy for these mixtures was midway between the 2HEP and 15HEP curves (Fig. 7a). However, replotting these data as a function of the surface occupancy of active (DRX) myosin (the product of myosin surface occupancy and predicted percent DRX (i.e., 100%-(predicted %IHM/SRX content)), Fig. 7b) demonstrates that the same relationship determines the velocity profiles for both the 2HEP and 15HEP constructs, as well as mixtures of the two. This suggests that the Velocity Index reflects the actin filament motility generated by active motors only, regardless of tail length and that IHM/SRX motors are mechanically silent, offering no resistance or internal load to the active motor population.

**Figure 7.**
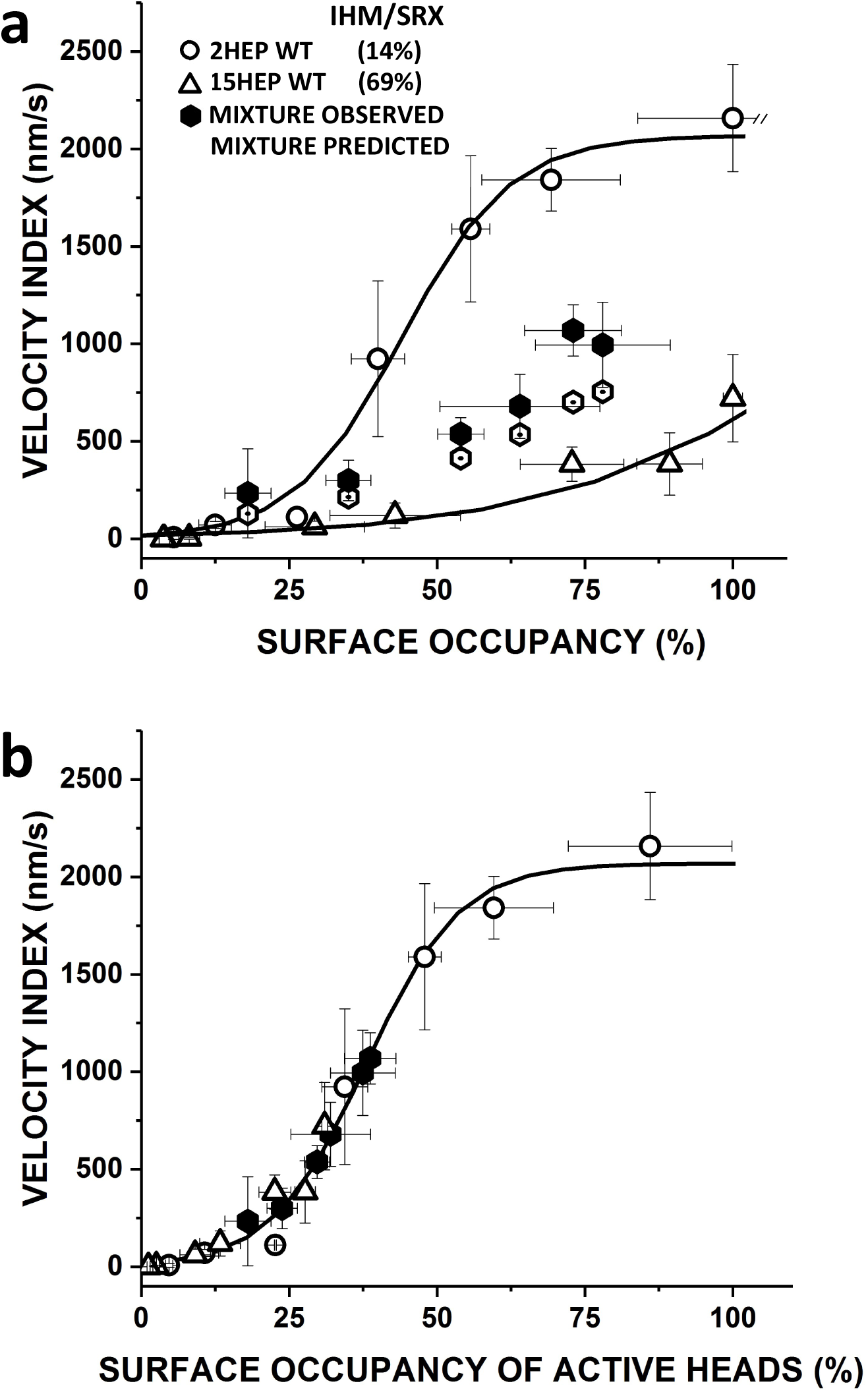
Influence of IHM/SRX Content on Motility of WT 2HEP:15HEP Mixture. a) The Velocity Index versus Surface Occupancy of WT 2HEP, 15HEP and a mixture of these two constructs. Addition of increasing amounts (0, 25, 50, 75, 100, 125 nM) of 15HEP WT (76% IHM/SRX) to a constant amount (25nM) of 2HEP WT (14% IHM/SRX) results in increasing Velocity Index values (filled octagons) that compare favorably to predicted values (open octagons; see Supplemental Table 1), supporting the model’s predictive nature, which assumes that a given construct has a defined IHM/SRX content and that the Velocity Index reflects actin filament motility generated by only active motors. b) Data in “a” were transformed to Velocity Index versus Surface Occupancy of Active Heads by replotting the Velocity Index data in “a” after calculating the surface occupancy of active (DRX) myosin heads from the product of observed total myosin surface occupancy and predicted percent DRX (i.e., 100%-(predicted %IHM/SRX content). The solid curve represents the 0% IHM/SRX curve (Supplemental Fig. 4). The 2HEP (open circles), 15HEP (open triangles), and observed mixture (closed hexagons) data, once transformed to represent only active motors, are well fit by the 0% IHM/SRX curve. Data points represent mean values ± SD of three experiments from separate protein preparations.

### dATP reduces the IHM/SRX population

deoxy ATP (dATP) has been shown to shift myosin out of the IHM/SRX state, while still supporting myosin’s biochemical and mechanical functions.^29^ Therefore, dATP should increase the Velocity Index for constructs with significant IHM/SRX content. To test this, dATP was exchanged for ATP in the motility assay with the WT 2HEP and 15HEP constructs at 100% myosin surface occupancy. Figure 8 shows no significant change in the WT 2HEP Velocity Index in the presence of dATP compared to that with ATP, (p = 0.94). In contrast, dATP increased the WT 15HEP Velocity Index by ∼50% (p = 0.01), which our modeling would predict is due to a ∼10% reduction in IHM/SRX (Fig. 7b). These results support the biochemical and motility data that suggest a low IHM/SRX population in the WT 2HEP construct. Whereas, dATP apparently reduces but does not eliminate the high IHM/SRX population in the WT 15HEP construct.

**Figure 8.**
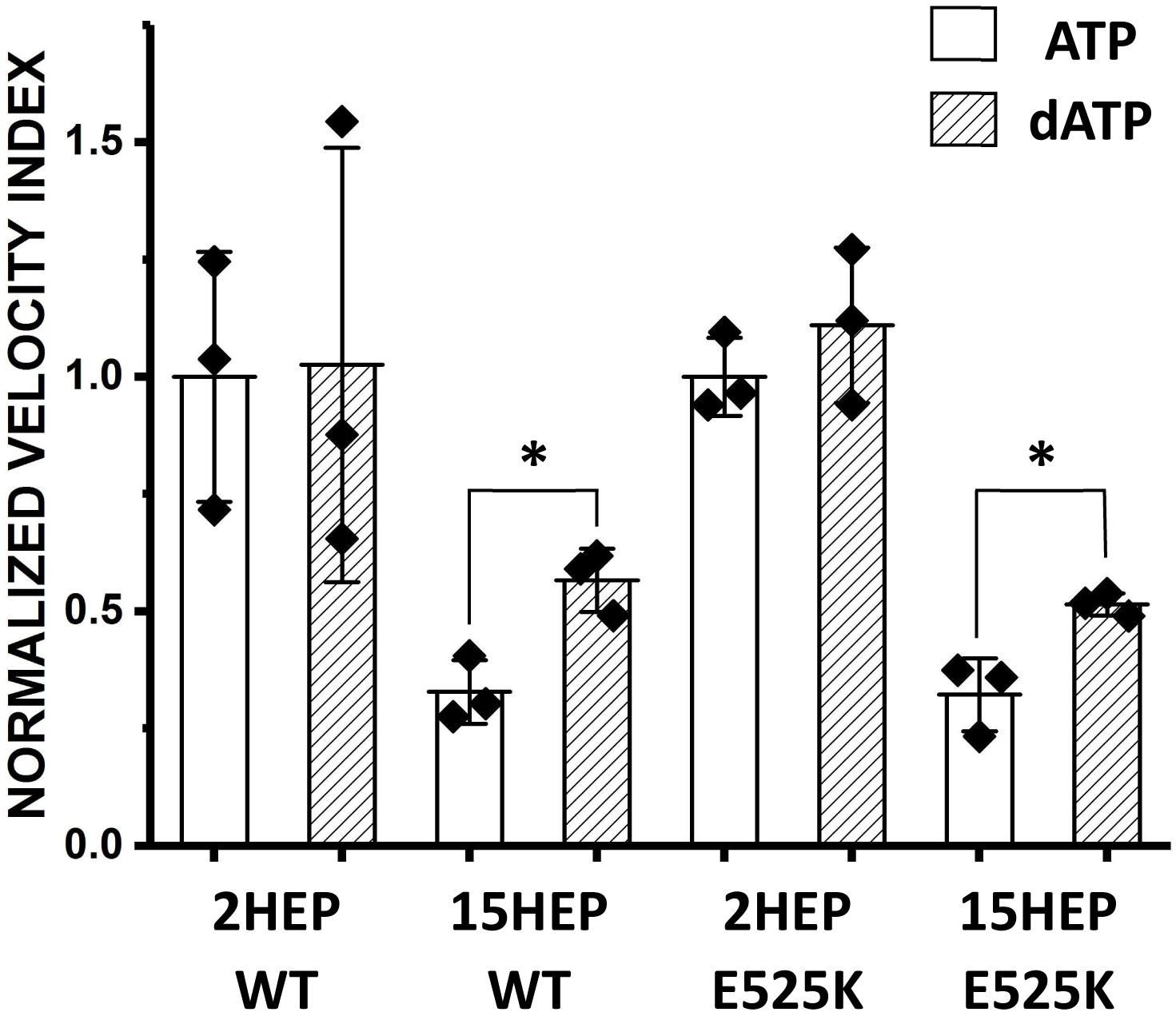
Deoxy-ATP (dATP) impact on IHM/SRX motors. The SRX inhibitor, dATP, was exchanged for ATP in the motility assay with the WT and E525K 2HEP and 15HEP constructs at 100% myosin surface occupancy. Data are normalized to the respective (i.e., WT and E525K) 2HEP mean Velocity Index data in the presence of ATP. * identifies significantly different than control (p=0.01). Data points represent mean values ± SD of three experiments from separate protein preparations.

### Impact of E525K mutation on ATPase activity and *in vitro* motility

#### Steady state and single turnover ATPase

Introduction of the E525K mutation in both the S1 and 2HEP constructs enhanced the steady state, actin-activated ATPase activity ∼3-fold as well as dramatically decreasing by ∼18-fold the actin concentration at which ATPase activity is one-half maximal (K_ATPase_) (Fig. 3a,b; Table 1), as reported previously^10^ for the S1. However, both the E525K 15HEP and 25HEP actin-activated ATPase activity appeared insensitive to the mutation compared to WT and continued to show reduced actin-activated ATPase activity compared to the E525K S1 and 2HEP constructs (Fig. 3; Table 1).

As for single ATP turnover measurements that were used to assess a construct’s fraction of SRX, the E525K mutation did not alter the S1 and 2HEP SRX percentage, which remained small (5-10%) across all KCl concentrations (Fig. 4a,b; Tables 2,3). The SRX rate constants were unchanged in the E525K S1 and 2HEP constructs, whereas the DRX rate constant was accelerated approximately 2-fold by the mutation at almost all KCl concentrations (Table 2). Compared to WT, the E525K mutation in both the 15HEP and 25HEP constructs stabilized the SRX state against changes in ionic strength (Fig. 4c,d; Tables 2,3). Notably, in the E525K 25HEP construct, the SRX rate constants were universally decreased approximately 2-fold, whereas the DRX rate constants were primarily unchanged, except at the higher salt concentrations (100 and 150 mM KCl) where they were accelerated approximately 3-fold (Table 2).

#### In vitro motility

The Velocity Index versus myosin surface occupancy relationship for the E525K short- (2HEP) and long-tailed (15HEP and 25HEP) constructs were compared to WT relations (Fig. 9). All three E525K constructs were not significantly different from their WT counterparts (2HEP, p = 0.75; 15HEP, p = 0.75; 25HEP, p = 0.87). Using our model to estimate the %IHM/SRX for the various E525K constructs (2HEP, 15HEP, 25HEP) (see above), the least squared error fits to the Velocity Index versus myosin surface occupancy relations (Fig. 9, dashed curves) suggest IHM/SRX percentages of 16±12% (E525K 2HEP), 62±6% (E525K 15HEP) and 71±4% (E525K 25HEP). Given the error in the predicted %IHM/SRX estimate for the various model fits (Suppl. Fig. 4), the predicted %IHM/SRX for all of the E525K mutants are no different than WT. Additionally, the lack of difference between WT and E525K 2HEP constructs, which do not contain a large IHM/SRX population (Fig. 4b), suggests that the mutation does not impact the motor’s inherent motion generating capacity under the unloaded conditions in the motility assay. In addition, the E525K mutation does not appear to further stabilize or alter the long-tailed 15HEP and 25HEP ability to adopt the IHM/SRX state at least for the low salt conditions used in the motility assay.

**Figure 9.**
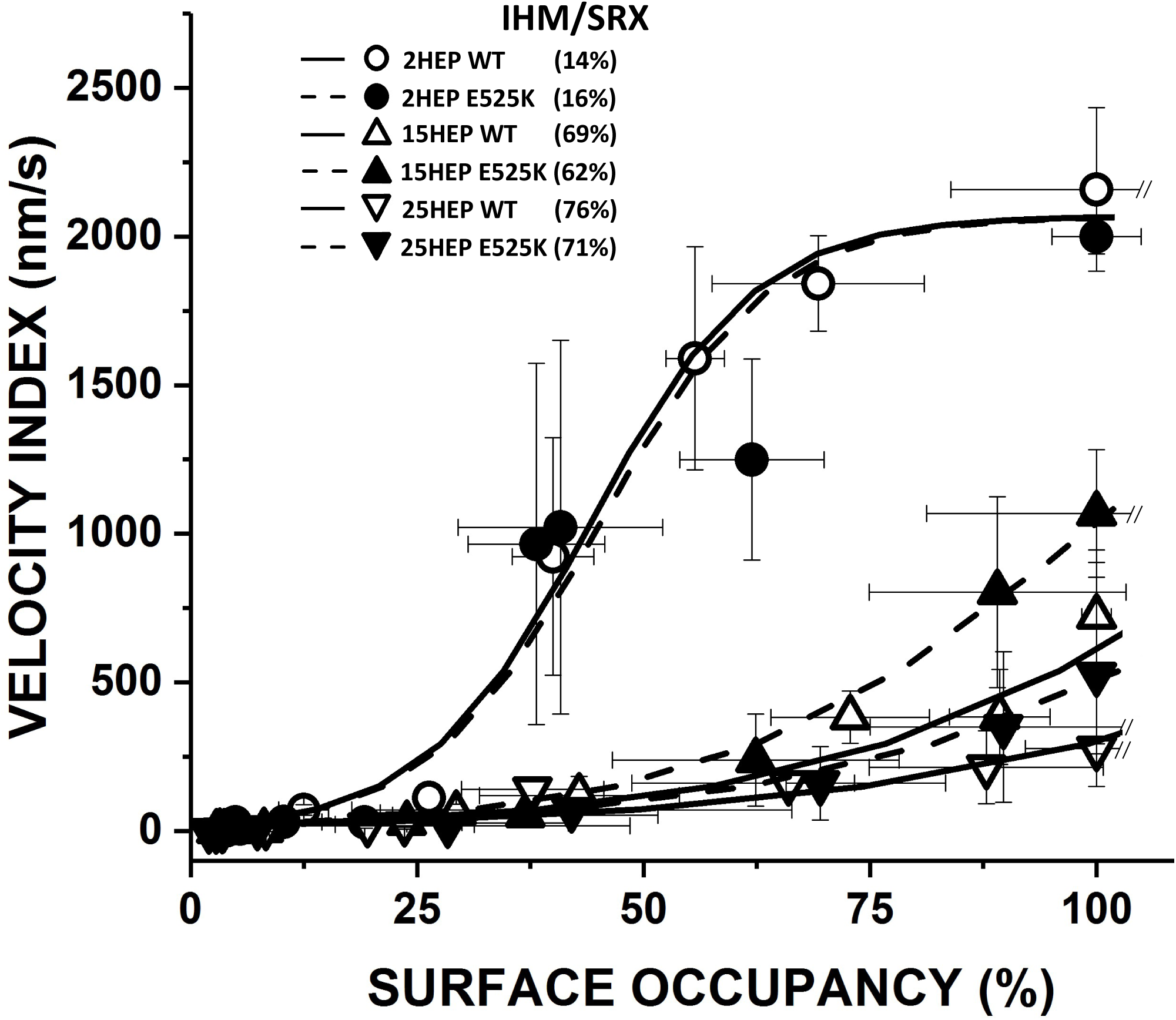
Impact of the E525K mutation on M2β motility. Velocity Index versus Surface Occupancy for WT and E525K 2HEP, 15HEP, and 25HEP constructs. WT data and fitted curves from Figure 6b. All three E525K constructs were not significantly different than their WT controls (2HEP, p = 0.75; 15HEP, p = 0.75; 25HEP, p = 0.87). Dashed curves are the least squared error model fits to the Velocity Index versus Surface Occupancy relations in order to estimate the %IHM/SRX for the various E525K constructs (see legend in figure). The predicted %IHM/SRX for all of the E525K mutants are no different than WT. All *in vitro* motility experiments performed in low salt (25 mM KCl). Data points are presented as mean ± SD of at least three experiments from separate protein preparations.

As with the WT constructs, we exchanged dATP for ATP in the motility assay for the E525K constructs to determine if in contrast to the WT 15HEP construct, the mutation may prevent the transition out of the IHM/SRX state due to the presence of dATP. As might be expected for the E525K 2HEP construct due to its low IHM/SRX percentage (Fig. 4b; Tables 2,3), dATP had no significant effect on the normalized Velocity Index (p = 0.38) (Fig. 8). In contrast, the presence of dATP produced an ∼40% increase in the normalized Velocity Index for the E525K 15HEP construct (p = 0.01) (Fig. 8), suggesting that additional motors are recruited out of the IHM/SRX state. Notably, the similar increase in Velocity Index for both the WT and E525K 15HEP due to dATP further suggests that the E525K mutation does not enhance the stability of the IHM/SRX state in the 15HEP construct at least under the low ionic strength conditions of the motility assay.

## Discussion

Hypertrophic (HCM) and Dilated Cardiomyopathy (DCM) are considered diseases of the sarcomere, as these myopathies involve mutations in β-cardiac myosin and other sarcomeric proteins critical to muscle function^33-36^. Although cardiac contractility in HCM and DCM patients differ dramatically, i.e., hyper-contractile^37^ and hypo-contractile^38^, respectively, the underlying myofilament-based mechanisms that dictate these contractile differences may be common. Specifically, cardiac muscle power generation is related to the number of force-generating myosin motors recruited upon activation and each motor’s intrinsic force and motion-generation while attached to the actin-thin filament (Fig. 1). Therefore, HCM- and DCM-associated mutations to myosin could impact either or both of these critical processes, with evidence in the literature for both^33,39^. Garnering much attention is the super-relaxed, auto-inhibited myosin state (SRX) that can be stabilized by intramolecular interactions between myosin’s two heads and its coiled-coil tail domain (i.e. interacting heads motif, IHM)^10,40-42^. Therefore, the proportion of myosin that adopt this auto-inhibited IHM/SRX state effectively determines the number of available force-generating myosin heads upon activation (Fig. 1). In fact, HCM-associated myosin mutations appear to disrupt the IHM/SRX state, increasing myosin head availability upon cardiac activation, which may explain the observed hyper-contractility^39^. In contrast to HCM, do DCM-associated β-cardiac myosin mutations further stabilize the IHM/SRX state, thus reducing the number of functionally available myosin heads, which would lead to cardiac hypo-contractility (Fig. 1d). In fact, we previously reported that the E525K DCM mutation, in expressed human β-cardiac myosin with a 15 heptad tail (15HEP), did stabilize the IHM structural state and thus the SRX biochemical state as well, resulting in a reduced actomyosin ATPase activity compared to wildtype (WT) myosin^10^. Here we extend these studies in both WT and E525K DCM mutant human β-cardiac myosin to better define the extent that the IHM/SRX state is dependent on the myosin tail length, and how the IHM/SRX state impacts both myosin’s hydrolytic and molecular mechanical function with and without the E525K mutation.

Here we show, as have others^9,43^, that myosin’s ability to adopt the IHM/SRX state, based on steady-state actomyosin ATPase and single ATP turnover experiments, requires both heads and at least 15 heptads of tail length. This tail length was reported originally for smooth muscle myosin; where adopting the IHM state is a regulatory mechanism for inhibiting this myosin motor^44^. Therefore, as expected, a double-headed β-cardiac myosin construct with a short 2 heptad tail (2HEP) is enzymatically indistinguishable from a single-headed myosin S1 construct, which is incapable of adopting the IHM state (Fig. 3a,b). Whereas, the WT 15HEP and longer-tailed 25HEP constructs, which can adopt the IHM/SRX state, both exhibit significantly reduced steady-state actomyosin ATPase activity compared to WT S1 and 2HEP (Fig. 3; Table 1). The IHM/SRX state in the long-tailed constructs is stabilized by intramolecular electrostatic interactions^8-11^, as evidenced by a reduced fraction of myosin in the IHM/SRX state with increasing ionic strength (i.e., 20mM to 150mM KCl) (Fig. 4c,d). Although the WT 25HEP construct was less sensitive to ionic strength (Fig. 4d), suggesting that increased tail length and presumably additional intramolecular interactions lead to a more stable IHM/SRX state in solution^45^.

Myosin is a mechanoenzyme, having both enzymatic and mechanical activities. Although these dual activities are generally correlated, the rate limiting steps for the enzymatic and mechanical cycles differ^20,46^. Therefore, the *in vitro* motility assay has become a standard for characterizing the motor’s motion and force-generating capabilities^5,20,26,27^, which we used here to define how the β-cardiac myosin tail impacts myosin mechanical function. Specifically, the actin filament Velocity Index vs. myosin surface occupancy relation in the motility assay should reflect both myosin’s intrinsic motion generation and the number of motors available to interact and propel the actin filament, both of which might be altered by a DCM mutation. The non-linear, Velocity Index dependence on surface occupancy (i.e., motor number) (Fig. 6) is due to β-cardiac myosin being a low duty ratio motor^46^, i.e., spending a small fraction of its hydrolytic cycle attached to actin and generating motion. If the IHM/SRX state dictates the number of available motors, then the shape of the Velocity Index vs. myosin surface occupancy relation should be sensitive to the IHM/SRX state stability. Therefore, we took advantage of the M2β construct C-terminal, EGFP tag both as a site-specific attachment strategy to the motility surface and as a fluorescence indicator of the total number of motors on the motility surface (i.e., active and inactive (IHM/SRX)) at any myosin concentration. Using an antiGFP nanobody attachment to the surface, myosin heads are positioned to interact properly with actin without surface interference (See Suppl. Fig. 2) and are free to transition between the inactive IHM/SRX and active DRX states.

Based on each construct’s EGFP fluorescence, we confirmed that the myosin surface occupancy (i.e., total number of attached motors) was the same for any double-headed construct (Fig. 5), which if not the case would have confounded the interpretation of Velocity Index differences. Interestingly, the maximum Velocity Indexes for the WT 15HEP and 25HEP constructs were 3-fold slower than the WT single-headed S1 and double-headed, short 2HEP construct (Fig. 6). Based on enzymatic characterization, these same long-tailed constructs (i.e., 15HEP and 25HEP) can adopt the IHM/SRX state (see Results). Therefore, is the reduced WT 15HEP and 25HEP Velocity Index due to a reduced number of available heads? In support of this scenario, substituting dATP, an SRX inhibitor^29^, for ATP in the motility assay, the WT 15HEP Velocity Index increased by ∼50% (Fig. 8), suggesting that dATP shifted the myosin state equilibrium out of the IHM/SRX state, effectively making more active myosin available on the motility surface.

With the Velocity Index vs. myosin surface occupancy sensitive to the number of active myosin on the motility surface, we developed a simple analytical model (see Results and Supplemental Methods) to predict the proportion of WT 15HEP and 25HEP motors in the IHM/SRX state. For this model, we assumed, based on single ATP turnover experiments (Fig. 4b), that the WT 2HEP Velocity Index vs. myosin surface occupancy relation was that of a myosin population with 16% in the IHM/SRX state. Additionally, we assumed that IHM/SRX myosins are inactive and do not contribute to actin movement, thus effectively reducing the number of active heads at any myosin surface occupancy. Based on the model, the WT 15HEP and 25HEP constructs are predicted to have 69% and 76% IHM/SRX state probability, respectively (Fig. 6). These model estimates are in relatively good agreement with the ∼85% IHM/SRX proportion observed in the single ATP turnover experiments for these same constructs. (Fig. 4). In support of the model assumption that the IHM/SRX state is mechanically silent, thus effectively reducing the number of available motors, we conducted a mixture experiment using WT 2HEP and 15HEP in various ratios. This experiment allowed us to vary the proportion of IHM/SRX myosin on the motility surface, given the predicted 14% and 69% IHM/SRX state probability for the two constructs, respectively. The agreement between the observed and predicted Velocity Index vs. myosin surface occupancy for the various WT mixtures (Fig. 7) further supports a mechanism by which the IHM/SRX state effectively limits the number of active motors that contribute to cardiac contractility.

With both the WT myosin construct enzymatic and motile properties characterized, we then used the same experimental approaches to determine if the E525K DCM mutation impacts these properties and specifically if the mutation further stabilizes the IHM/SRX state, as suggested by our earlier study using only the S1 and 15HEP constructs. The E525K^10,47^ DCM-associated mutation is located in the myosin lower 50 kDa domain that is involved in electrostatic intramolecular interactions between the "blocked" head and the S2 domain that stabilizes the IHM state^36,48^. With the E525K mutation introducing a positive charge, electrostatic interactions could be strengthened and thus further stabilize the IHM/SRX state.

Interestingly, as we reported previously^10^, the E525K S1 showed ∼3-fold enhanced actomyosin ATPase activity relative to WT (Fig. 3a; Table 1). As expected, this enhanced ATPase activity was also observed in the E525K 2HEP (Fig. 3a,b; Table 1). However, no such ATPase enhancement was observed for the E525K 15HEP as reported previously^10^, and similarly for the E525K 25HEP (Fig. 3c,d; Table 1). With the myosin heads being identical for all of the mutant constructs, the lack of any effect of the E525K mutation on the 15HEP and 25HEP ATPase activities suggests that these long-tailed construct’s ability to adopt the IHM/SRX state may effectively mask the enhanced ATPase activities of the individual mutated heads. With respect to *in vitro* motility, there were no statistical differences in the Velocity Index vs. myosin surface occupancy relations for any of the double-headed E525K constructs compared to their WT controls (Fig. 9). The fact that the ∼3-fold greater actomyosin ATPase activity for the E525K 2HEP did not translate into a faster Velocity Index may reflect differences in the rate limiting steps for solution-based ATP hydrolysis versus surface-attached motion generation^20,46^.

If no differences in enzymatic and motile activities are apparent in the E525K 15HEP and 25 HEP constructs compared to WT, then how is the hypo-contractile DCM phenotype possible? Both the actomyosin ATPase and motility assay measurements are performed at low ionic strength (25mM KCl) to maximize actomyosin interactions in these reductionist assays. Whereas *in vivo*, the close proximity between the actin-thin and myosin-thick filaments within the muscle sarcomere make their effective concentrations infinite and thus overcome the electrostatic shielding effects of the high ionic strength in muscle. Based on the single ATP turnover data, at 20mM KCl, the IHM/SRX fraction (∼85%) for the E525K 15HEP and 25HEP are unchanged compared to WT. If the IHM/SRX state is the dominant factor dictating the actomyosin ATPase activity and actin filament motility *in vitro*, then it’s not surprising that the Velocity Indices were the same at low ionic strength for both E525K and WT long-tailed constructs. However, single ATP turnover measurements can be performed over a range of ionic strengths (Fig. 4; Table 3) and interestingly, the IHM/SRX probabilities for both E525K 15HEP and 25HEP constructs remained high and were insensitive to ionic strength up to 150mM KCl. These data suggest that the E525K DCM-associated mutation does stabilize the IHM/SRX state. Therefore, if both the steady-state actomyosin ATPase activity and *in vitro* motility assay could be performed at physiological ionic strengths, where the WT IHM/SRX state is destabilized, one might expect significantly reduced hydrolytic and motile behaviors from the DCM mutant myosin, which might explain the cardiac hypo-contractility in DCM patients. In the future, power generation, a more relevant physiological measure of cardiac muscle contractility, can be assessed at higher physiological ionic strengths in the optical trap^6,19,49^, whereby an actin filament can be brought into contact with a mutant myosin ensemble regardless of the ionic strength. By this approach the true impact of DCM-associated mutations on human β-cardiac myosin can be defined.

Given myosin’s hydrolytic and mechanical activities, which are each multi-step processes, there is no a priori reason to assume that every HCM or DCM mutation will impact these processes identically. Whether or not alterations in the IHM/SRX stability are the governing factor that dictates the overall biochemical or motile properties for every mutant myosin, which may be the case for the E525K mutation reported here, is yet to be established. DCM remains a formidable challenge due to its heterogeneous nature and the lack of a unifying molecular mechanism for monogenetic cases. Extending these findings to clinical settings and studying a broader spectrum of DCM mutations compared to E525K will hopefully provide more comprehensive insights into DCM’s underlying mechanisms.

## Acknowledgments

The authors thank Guy Kennedy from the Instrumentation and Model Facility at the University of Vermont for his microscopy expertise, Samantha Previs for her technical expertise, and the entire Warshaw lab for critical discussions. This work was supported by funds from the American Heart Association (23PRE1013839 to S.D.-M.), National Institutes of Health (HL150953 and HL163585 to D.M.W., C.M.Y., and S.S.), National Science Foundation (GRFP2237827 to D.V.), and a generous gift from Arnold and Mariel Goran to D.M.W.

