## Supplemental Material for "Tail Length and E525K Dilated Cardiomyopathy Mutant Alter Human β-Cardiac Myosin Super-Relaxed State"

**Model for impact of percent IHM/SRX on the Velocity Index versus Surface Occupancy.**

Our simple analytical model for the effect of IHM/SRX myosin on the Velocity Index versus Surface Occupancy relation assumes that the WT 2HEP relation is that of a myosin population with a 16% IHM/SRX content, based on the single ATP turnover data at 20 mM KCl (Fig. 4b). The WT 2HEP Velocity Index versus Surface Occupancy data were then fit to a modified Hill dose-response equation of the form: $= A1 + \frac{A2-A1}{1+1^{\ (C-x)p}}$ , using the Levenberg–Marquardt algorithm (OriginLab Corporation, Northampton, MA, USA), where $A1$ and $A2$ are lower and upper asymptotes of 0 and 2068±105 (nm/s), respectively, $C$ = 44±2% occupancy (i.e., an $EC_{50}$), and with a curve steepness factor, p = 0.05±0.01.

The model then uses the WT 2HEP Velocity Index versus Surface Occupancy does-response fit to generate a curve for a myosin that has 0%IHM/SRX content by simply shifting the WT 2HEP fit leftward along the $OCCUPANCY$ axis (i.e., X-axis) by dividing each $OCCUPANCY_{WT\ 2HEP}$ by 1.16 (i.e., the assumed 16%IHM/SRX). From this 0%IHM/SRX curve, a new family of curves for 0% to 100%IHM/SRX content was generated (Suppl. Fig. 4b) by simply shifting the 0%IHM/SRX curve rightward after mathematically rescaling the total myosin surface occupancy to be effectively equivalent to the % active myosin as follows:

$$OCCUPANCY_{new}(\%\text{IHM/SRX}) = \frac{OCCUPANCY_{0\%IHM/SRX}}{1-((\%IHM/SRX)/100))},$$

where $OCCUPANCY_{new}$(%IHM/SRX) denotes the adjusted total myosin surface occupancy for a myosin with a given IHM/SRX percentage, and $OCCUPANCY_{0\%IHM/SRX}$ is the total myosin surface occupancy associated with the model 0%IHM/SRX curve.

The predicted family of curves were then used to estimate the %IHM/SRX content for the various constructs. The observed Velocity Index versus Surface Occupancy data for a given construct were then iteratively fit to the family of curves in Suppl. Fig. 4b and a COST function generated by calculating the least sum of the squares of the residuals for the experimental data fit to each of the predicted family curves. The minimum of the COST function (Suppl. Fig. 4a, inset) was then taken as the best fit for the predicted %IHM/SRX content.

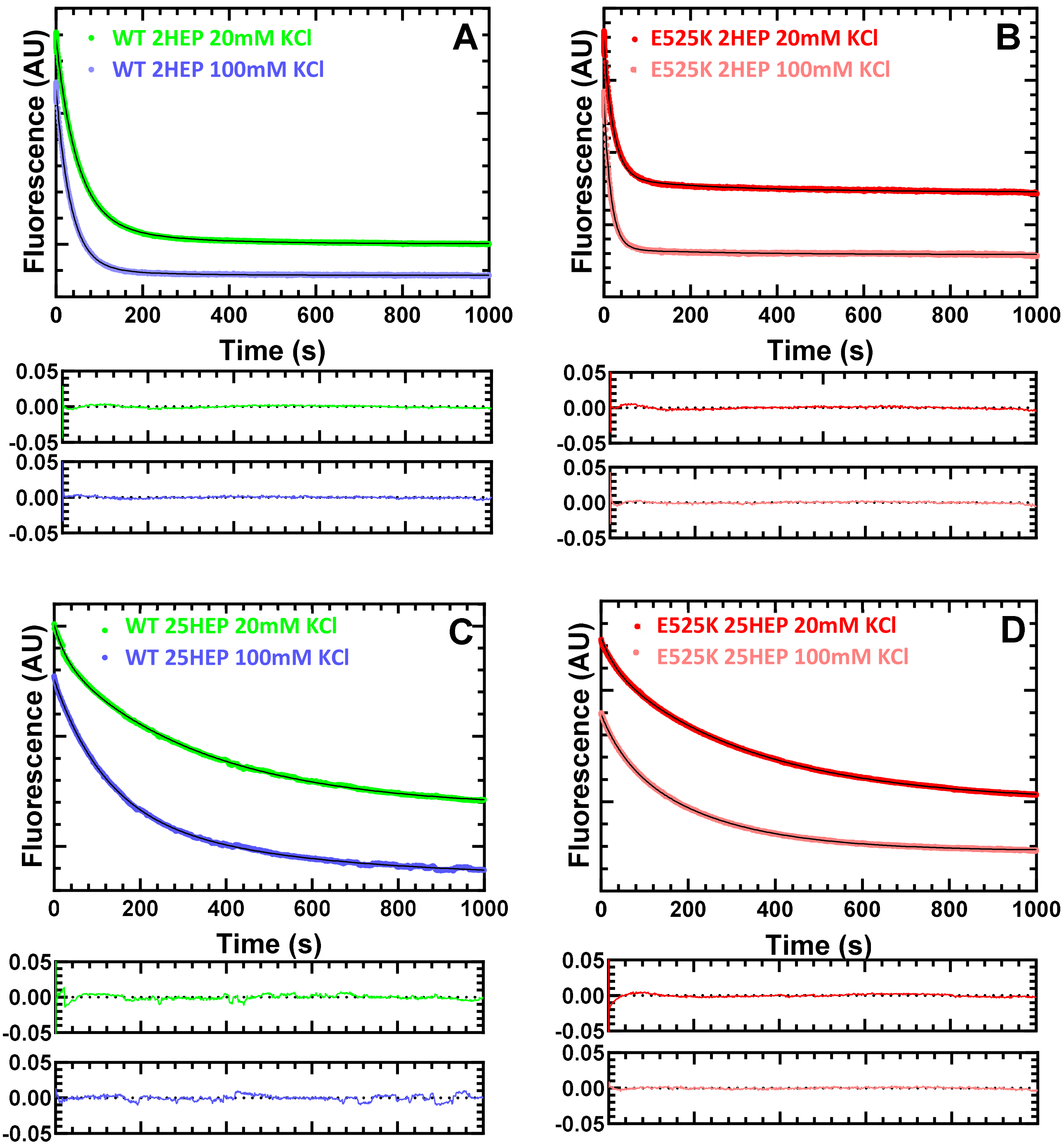


**Supplemental Figure 1. Single ATP turnover fluorescence decays.** Examples of single mantATP turnover reactions for WT and E525K 2HEP, and 25HEP constructs as a function of ionic strength (20mM and 100mM KCl). Representative fluorescence decay fits (upper graph) using two-exponentials (thin solid black line) and their respective residuals (lower pair of graphs) are presented. Relative amplitudes of the fast and slow phase exponential fit rate constants determined the fraction of myosin in the DRX and SRX states, respectively. See main text Table 2 for summary of values.

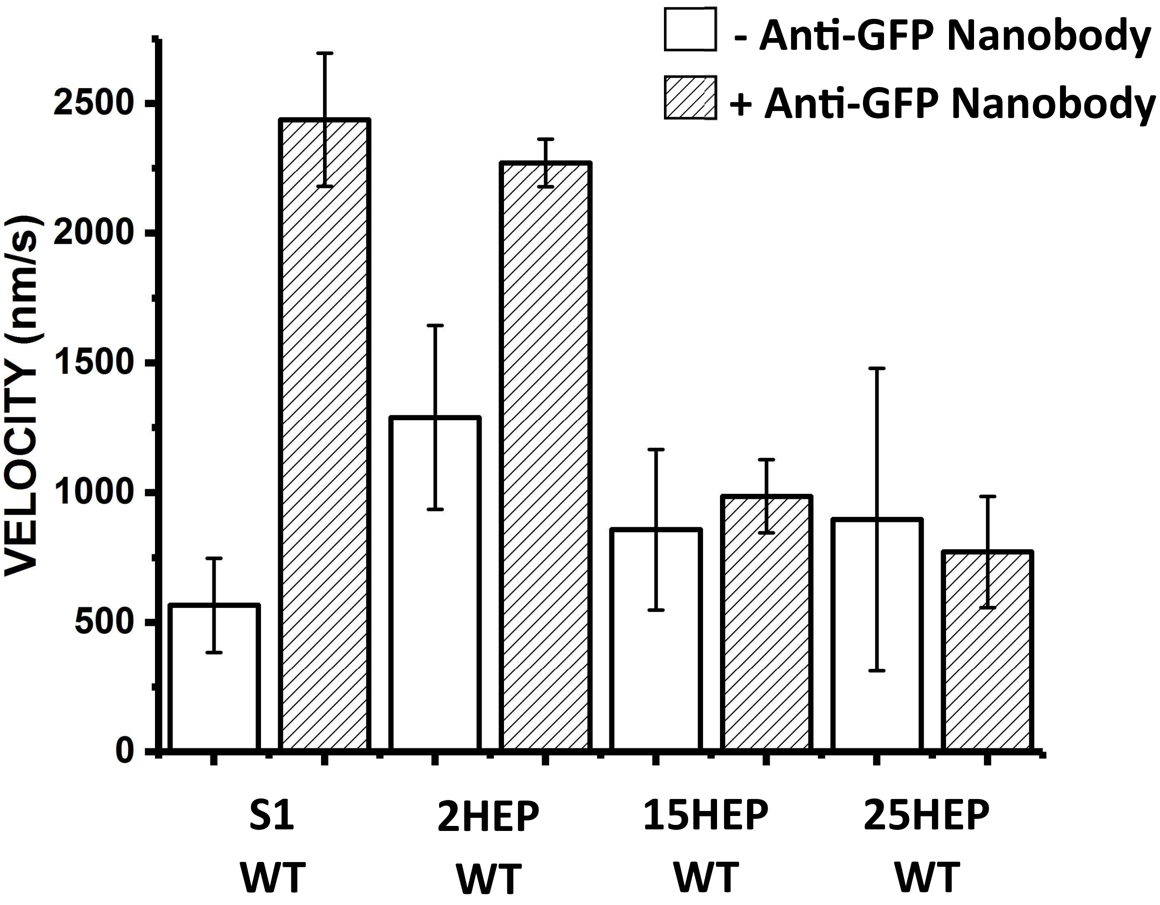


**Supplemental Figure 2. M2β motility surface attachment strategy.** The use of an anti-GFP nanobody to attach myosin constructs to the motility surface improves velocity of the S1 and 2HEP WT constructs, which are incapable of adopting the IHM/SRX state, by at least a factor of 2. This result suggests that the nanobody attachment strategy allows myosin heads to interact properly with actin without surface interference. Data represent mean values ± SD from three independent protein preparations.

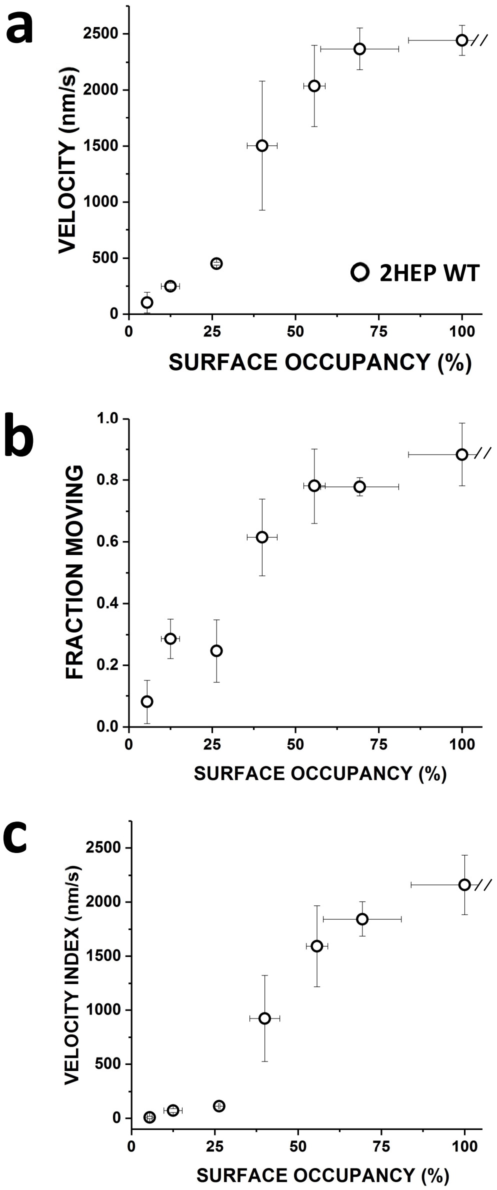


**Supplemental Figure 3. Velocity times Fraction Motile = Velocity Index.** The Velocity Index offers a more complete characterization of the motility behavior by multiplying velocity data by the fraction of moving filaments, which can be reduced at limiting myosin surface densities. a) 2HEP WT Velocity increases in proportion to the surface occupancy. b) 2HEP WT Fraction Moving is reduced at low myosin surface occupancy where the numbers of active motors becoming limiting. c) 2HEP WT Velocity Index = (Velocity) x (Fraction Moving) using data in "a" and "b". Data points represent mean values ± SD of three experiments from separate protein preparations.

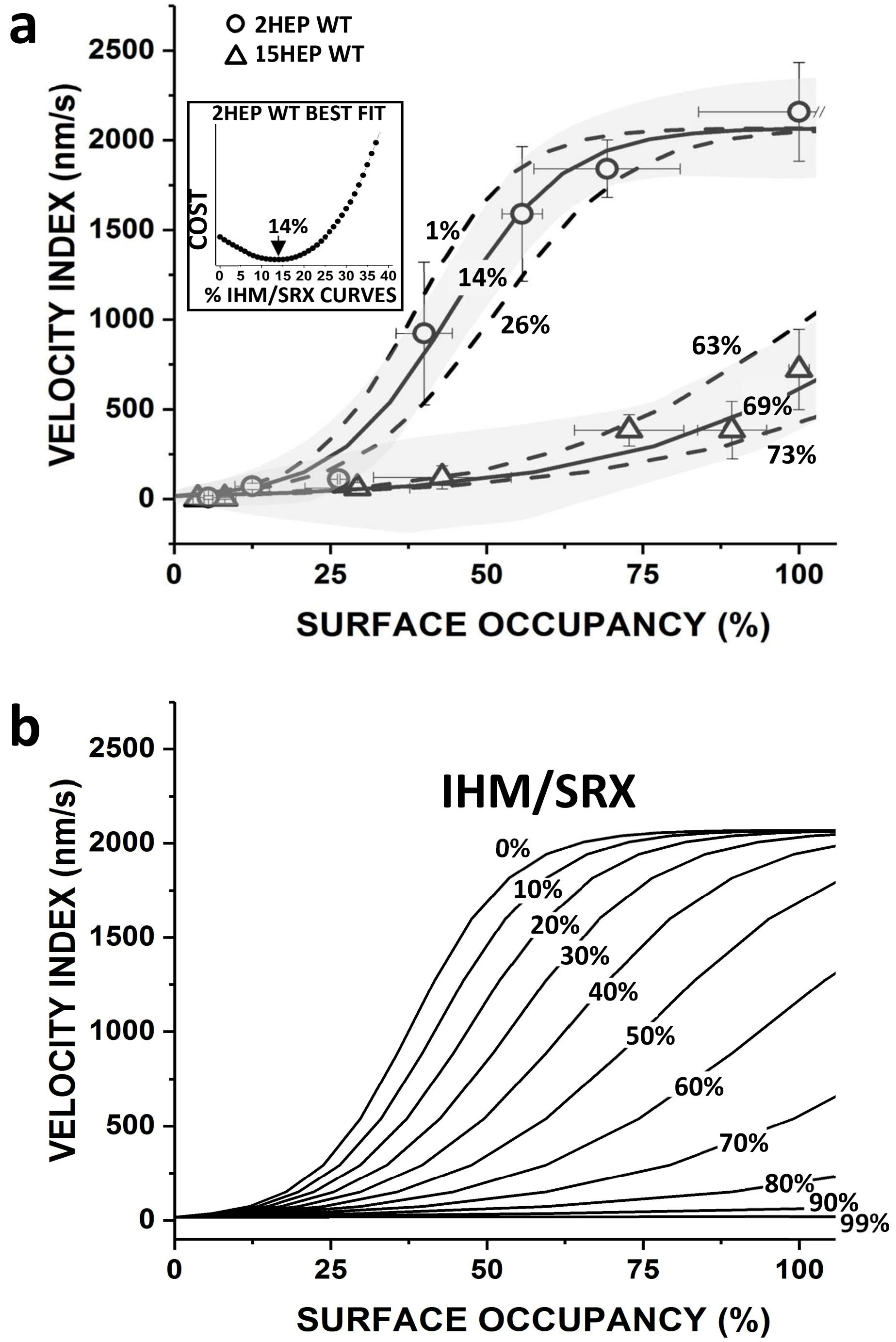


**Supplemental Figure 4. Model of the %IHM/SRX impact on M2β Velocity Index.** a) Velocity Index versus Surface Occupancy data for 2HEP WT were fit to the modified Hill equation (solid line) (Supplemental Methods) and used to represent an M2β population with 16% SRX that allowed a family of curves to be generated for hypothetical IHM/SRX percentages ranging from 0 to 100% in "b". The 95% confidence limits (gray shaded area) for the 2HEP WT fit were used to estimate the error in the model predicted %IHM/SRX for a given construct. All fits were performed by calculating the minimum COST function from the least squared error analysis (example for 2HEP WT, inset). For 2HEP WT the model best fit returns a best fit of 14% IHM/SRX with an estimate error of ±13% IHM/SRX. Velocity Index versus Surface Occupancy data for 15HEP WT with model predicted 69% IHM/SRX curve and 95% confidence limits (gray shaded area) error estimates of ±6% IHM/SRX. Predicted %IHM/SRX and the errors in the estimate for all constructs were as follows: WT: 2HEP (14±13%), 15HEP (69±6%), 25HEP (76±4%); E525K: 2HEP (16±12%), 15HEP (62±6%), 25HEP (71±4%). b) Family of curves for hypothetical %IHM/SRX ranging from 0 to 100% based on the original WT 2HEP modified Hill dose-response curve in "a", which assumed a 16%IHM/SRX (see Supplemental Methods for details).
